# Diverse ancestral brainstem noradrenergic neuron activity across species and biological factors

**DOI:** 10.1101/2024.10.14.618224

**Authors:** Michael A. Kelberman, Ellen Rodberg, Brandon R. Munn, Ehsan Arabzadeh, Chloe J. Bair-Marshall, Craig W. Berridge, Esther Berrocoso, Vincent Breton-Provencher, Daniel J. Chandler, Alicia Che, Oscar Davy, David M. Devilbiss, Anthony M. Downs, Gabrielle Drummond, Roman Dvorkin, Zeinab Fazlali, Robert C. Froemke, Erin Glennon, Joshua I. Gold, Hiroki Ito, Xiaolong Jiang, Joshua P. Johansen, Alfred P. Kaye, Jenny R. Kim, Chao-Cheng Kuo, Rong-Jian Liu, Yang Liu, Meritxell Llorca-Torralba, Jordan G. McCall, Zoe A. McElligott, Andrew M. McKinney, Cristina Miguelez, Ming-Yuan Min, Alexandra C. Nowlan, Mohsen Omrani, Gina R. Poe, Anthony Edward Pickering, Yadollah Ranjbar-Slamloo, Jone Razquin, Charles Rodenkirch, Anna C. Sales, Rath Satyasambit, Stephen D. Shea, Mriganka Sur, John Arthur Tkaczynski, Sonia Torres-Sanchez, Akira Uematsu, Chayla R. Vazquez, Amelien Vreven, Qi Wang, Barry D Waterhouse, Hsiu-Wen Yang, Jen-Hau Yang, Liping Zhao, Ioannis S. Zouridis, James M. Shine, David Weinshenker, Elena Vazey, Nelson K. Totah

**Author notes:** Denotes equal contribution (Michael A. Kelberman, Ellen Rodberg).

## Abstract

The brainstem cell group, locus coeruleus (LC), is present across vertebrates and influences cardiorespiratory, metabolic, immune, and cognitive functions by activating in two putatively distinct firing patterns. Yet, the degree to which the LC firing rates and patterns are homogenous across species has never been assessed due to inherently limited sample sizes. To remedy this, we pooled cross-species data from 20 laboratories to show that firing rates differ across species and are modulated by sex, age, and type of *in vitro* or *in vivo* preparation. Contrary to the prevailing dual-mode firing pattern schema, we observed patterns spread across a low-dimensional manifold, with subregions enriched for specific biological factors and neurodegenerative disease models. Our findings show considerable diversity in an ancestral vertebrate neuromodulatory system.

## Main Text

The central nervous system-wide projections of the brainstem noradrenergic cell group, locus coeruleus (LC), have been retained across the vertebrate phylogenetic tree (*1*). A core function of the LC – mobilization of cardio-respiratory, metabolic, and sensory-motor-cognitive resources in response to environmental change – has also been conserved across vertebrates (*2–7*). This function is thought to be implemented by two modes of activity (tonic or phasic) and, while this dual-mode terminology has been widely applied to human neuroimaging, cognitive psychology studies, and computational models of cognition and learning, we lack a cross-species cellular level definition of these activity modes (*8–10*).

Based on the evolutionary conservation of LC structure and function, we hypothesized that the firing modes of LC neurons have been conserved across vertebrates to fulfill a fundamental functional role. Consider the analogy of breathing, which is fundamentally the same function across mammals. Breathing in mammals, whether the individuals are male or female – young or old – asleep or awake – is driven by homologous brainstem activity patterns in the pre-Bötzinger nucleus (*11*) regardless of species or these biological factors. The same may be the case for the LC and its resource mobilization function.

To date, this hypothesis has never been tested due to the small sample sizes of individual studies. Typically, only a single neuron is recorded in a brain slice and, in anesthetized or awake animals, only one or two LC neurons are accessible in extracellular recordings (*12–15*), which can in principle record activity of many neurons (“single units”) simultaneously but have an extremely limited yield in the brainstem due to physical instability of the electrode-tissue interface caused by cardiac, respiratory, and spinal movement (*16*).

The LC, via its ascending brain-wide projections, is implicated in cognitive deficits observed in attention deficit and hyperactivity disorder, schizophrenia, and neurodegenerative diseases (*17, 18*) as well as psychological and physical symptoms of post-traumatic stress disorder, anxiety disorder, and symptoms of opioid withdrawal (*19–21*). Moreover, via descending projections to the autonomic nervous system and the spinal cord, the LC influences peripheral analgesia, cardiovascular disease, immune dysfunction, metabolic disorders, and obesity (*22–27*). The relevance of LC to such a broad range of human health and disease conditions underscores the importance of confirming or refuting whether LC activity varies across species, or across other factors such as sex, age, and experimental preparation.

To remedy this issue, we compiled spontaneously occurring spike trains of LC neurons collected across 20 laboratories (**Table S1**). These data included mice and rats of both sexes, as well as male *Macaca mulatta*, a non-human primate (NHP). Animals spanned relatively young (adolescent) to adult and late age. Spike discharges were recorded either in tissue slices (intracellular), intact animals under anesthesia of various types, or during wakefulness.

Additionally, some LC neurons or animals were genetically modified for optogenetic manipulation and, while recordings during optogenetic stimulation were not included in the data set, the genetic modification was considered as a factor that could influence firing rate. Finally, a subset of mice was pharmacologically manipulated or bred to express disease-specific genotypes. Laboratories providing extracellular recordings were asked to include only single units (henceforth referred to as “neurons”). However, in the interest of applying similar criteria to define single units across laboratories, we performed a quality control sweep of the 2,944 neurons, which yielded 1,855 and 1,708 neurons for firing rate and pattern analysis, respectively (see Supplementary Materials, *Quality control of single unit recordings across laboratories*).

Our goals were to compare neuronal activity levels (firing rate) and to describe tonic and phasic firing modes across species and across a range of intrinsic and extrinsic factors. To analyze neuronal activity levels, we analyzed time-averaged firing rate using negative binomial regression (a marginal model) to explain the individual and combined effects of species, sex, age, genotype, and brain state or experimental preparation type (e.g., awake, anesthesia of various types, or *ex vivo* slice recordings). There was a lack of substantial variation in firing rates between laboratories (**Figure S1**). We confirmed this by calculating an intraclass correlation value, which ascertains whether firing rates from the same laboratory were more similar than those obtained by other laboratories. The intraclass correlation value was 0.48, which suggests a low to moderate dependence of firing rate on laboratory identity. Including laboratory identity as a factor affecting firing rate (e.g., in a hierarchical model) was not possible because the low variability across laboratories prevented a hierarchical model from fitting the data. Given the limitations of the hierarchical model, we chose to use the marginal model. We examined the main effects of individual factors and considered interactions between these factors (**Table S2**). Some factors, such as animal strain, were not included in the model (see Supplementary Materials, *Factors excluded from the model of firing rates*). Effects, when significant, are reported as effect sizes in firing rate (spikes/sec) with 95% confidence intervals (CIs) to provide an intuitive quantification of how much firing rate differs between factors (*28–30*). Importantly, small effect sizes of even 0.5 spikes/sec may carry considerable relevance to brain function.

First, consider that the baseline discharge of LC neurons is already low (typically ranging from 0.5 to 5 spikes/sec) and that a relative change of 0.5 spikes/sec is a large difference in this range of activity. Second, our rudimentary understanding of how LC firing rate alters nervous system function largely precludes making a judgement regarding what effect size may (or may not) have consequences.

### LC firing rate is not conserved across model species and is influenced by brain state and sex

We start from the premise that LC firing rate is similar across model species due to the conservation of LC function across phylogeny. Our analysis showed a significant main effect of species (p<0.0001; **Figure 1A**; **Table S2**). Neurons recorded from mice had higher firing rates compared to rats (p<0.0001) and NHPs (p=0.0001), but rats and NHPs did not significantly differ from each other (see post-hoc comparisons in **Table S3**). The effect of species on LC activity interacted with the type of experimental preparation defining the state of the animal (p=0.001, **Figure 1B-F, Table S3**). The mouse LC is more active relative to the rat LC under all tested anesthetics and during *ex vivo* (brain slice) recordings (**Figure 1B-E, Table S3**). However, in the awake state, mouse LC neurons were less active compared to rats (p=0.0036) and NHPs (but to a lesser extent given the effect size 95% CIs overlap with no difference between means and p=0.0491) (**Figure 1F, Table S3**).

**Fig. 1.**
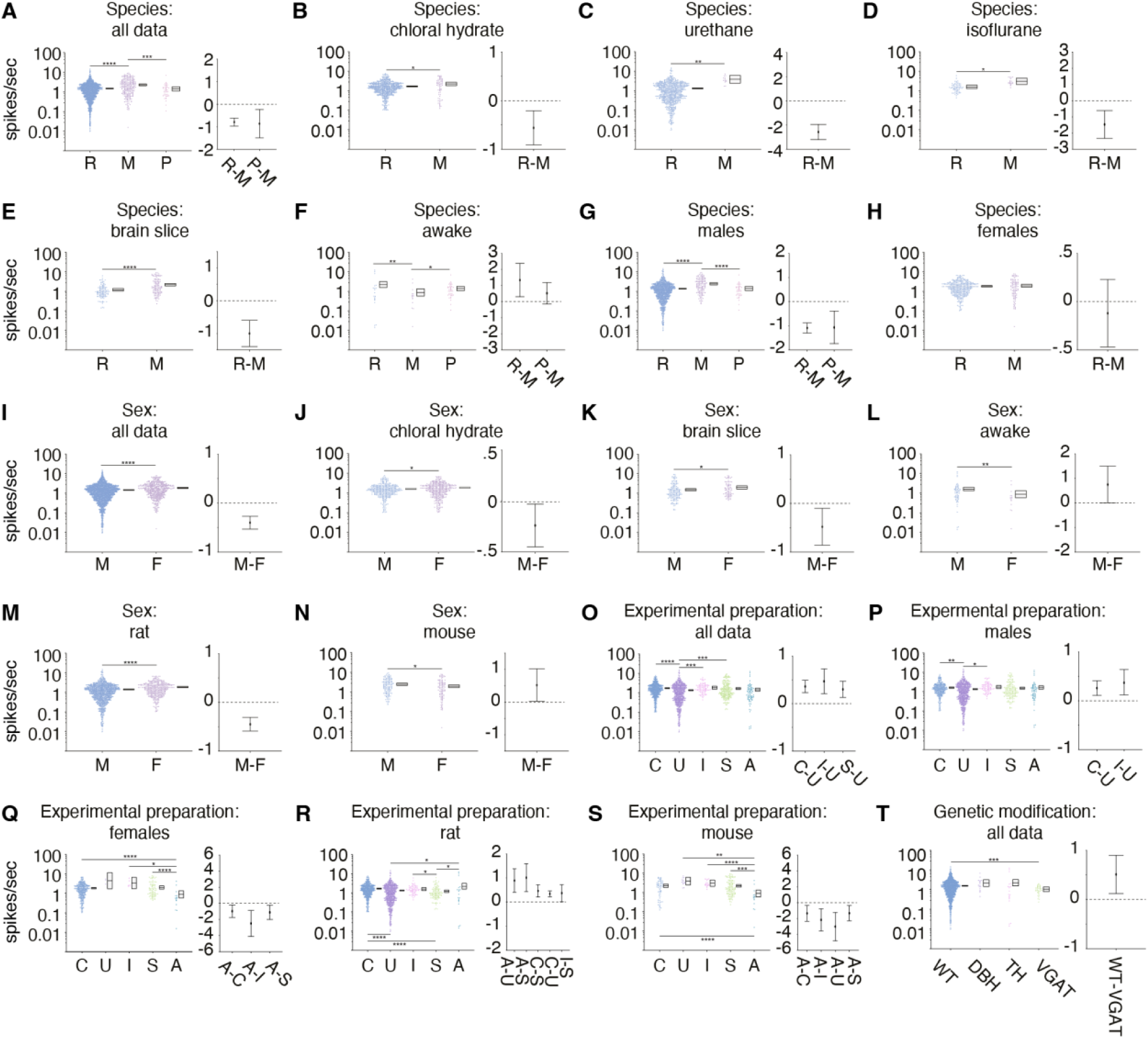
Dependence of LC firing rate based on interactions with intrinsic and extrinsic biological factors. All panels show the firing rate on a logarithmically-scaled y-axis on the left. The individual points are neurons. The sample size (N) can be found in table S1. The box next to the points shows the SEM of the firing rate. Asterisks indicate significant differences in firing rate according to the negative binomial regression model (*p < 0.05, **p < 0.01, ***p < 0.001, ****p < 0.0001). On the right, the effect size (i.e., difference between group means) with 95% confidence intervals (CIs) are shown for groups with significant differences in firing rate according to the negative binomial regression model. An effect is apparent if the 95% CI does not cross zero (dotted line). The effect sizes are shown in firing rate (spikes per second). (**A**) There was a main effect of species on LC firing rates, with mouse LC neurons displaying higher activity compared to both rats and non-human primates. (**B – E**) The effect of species also depended on the state of the animal. Abbreviations: R – rat, M – mouse. Mouse LC neurons were more active when recorded under chloral hydrate (**B**), urethane (**C**), isoflurane (**D**), and in slice (**E**). In awake animals, this effect was reversed, with rat and NPHs displaying higher firing rates compared to mice (**F**). (**G – H**) There was also a separate species and sex interaction. (**G**) Male rats and NPHs had lower firing rates compared to mice. (**H**) There was no difference between rat and mouse LC firing rates when comparing females, while we had no NHP data for comparison **I**) Sex had a main effect on firing rate. LC neurons recorded from female animals had a higher firing rate compared to males. . (**J-L**) Sex and brain state (experimental preparation) interacted. Female animals recorded under chloral hydrate (**J**) and in slice (**K**) have a higher firing rate than males. (**L**) Contrastingly, in male animals, LC firing rates were higher than females in the awake state. (**N -M**) Sex and species interacted to effect firing rate. In rats, females LC firing rates were higher than in males. (**M**) This effect was reversed in mice, where male LC neurons were more active than female LC neurons. (**O**) We observed a main effect of brain state on LC firing rates which revealed firing rates recorded under urethane anesthesia were lower than those recorded under chloral hydrate, isoflurane, and in slice. (**P-Q**) Brain state interacted with sex. (**P**) LC firing rates under urethane anesthesia were lower than chloral hydrate anesthesia and isoflurane anesthesia in male animals. (**Q**) In female animals, firing rate was lower in awake animals compared to chloral hydrate or isoflurane anesthesia and to slice. (**R-S**) Brain state interacted with species (only rodents could be tested due to the lack of NHP anesthetized recordings). (**R**) In rats, LC firing rates in the awake state were higher compared to those recorded under urethane anesthesia or in slice preparations. In addition, firing rates under chloral hydrate anesthesia were higher than in slice and urethane. Finally, firing rates under isoflurane anesthesia were higher than those recorded in slice preparations. (**S**) In mice, LC firing rates recorded in the awake state were lower than in all other states. (**T**) There was also a main effect of genetic modification (Cre expression) on LC firing rates, with VGAT-Cre animals demonstrating a lower firing rate when compared to WT animals. Abbreviations: R – rat, M – mouse, P – NHP, M – male, F – female, C – chloral hydrate, U – urethane, I – isoflurane, S – slice, A – awake, WT – wild-type, DBH – dopamine beta-hydroxylase, TH - tyrosine hydroxylase, VGAT - vesicular GABA transporter. *p < 0.05, **p < 0.01 ***p < 0.001, ****p < 0.0001.

The effect of species also interacted with sex (p=0.0103; **Figure 1G & H, Table S2**). The higher firing rate in mice relative to rats and NHPs was observed specifically in males (p<0.0001 vs rats; p<0.0001 vs NHPs) (**Figure 1G, Table S3**). The effect sizes between male mice versus male rats and male NHPs were 1.09 and 1.07 spikes/sec, respectively. On the other hand, in females, there were no species-specific differences between mice and rats (**Figure 1H, Table S3**). We did not have data from female NHPs for comparison. It has long been assumed that LC firing rate is conserved across model species, yet our findings show species-specific LC firing rate. Intriguingly, these differences depended on sex and animal state, which led us to directly examine sex and state differences in LC firing rate.

### Sex impacts LC firing rates in opposing directions in rats and mice

Biological sex is associated with differences in autonomic control, the stress response, sleep, and cognition, as well as symptoms of various clinical disorders (*31–36*). These sex differences could be associated with sex-specific LC firing rates. Our analysis revealed a main effect of sex on firing rate (p<0.0001; **Figure 1I**, **Table S2**). We found that the female LC was more active than male LC with an effect size of 0.4 spikes/sec. However, there was also an independent interaction between sex and brain state (p=0.0044; **Figure 1J-L**, **Table S2**). Specifically, the female LC was only more active when recorded under chloral hydrate (p=0.031) or from brain slice recordings (p=0.024). On the other hand, in awake animals, the female LC was less active compared to the male LC (p=0.004; **Figure 1J-L, Table S3**). The effect of sex also depended on species (p=0.0103; **Figure 1M & N, Table S2**). In rats, the female LC is more active than male LC (p<0.0001; **Figure 1M**, **Table S3**), whereas in mice the opposite was observed (p=0.034; **Figure 3N**, **Table S3**). Thus, LC firing rate varies by sex, but species and brain state modify the effect of sex.

### Firing rate in the awake state is increased in rats but decreased in mice

Given the technical challenges of targeting the LC with recording electrodes and sustaining recording signal quality, a substantial basis of our understanding of LC activity has been informed by recordings in anesthetized animals or in brain slices. The pharmacological manipulations (for anesthesia) or afferentation of the LC (for brain slice recordings) could entail a distinct brain state relative to awake animals. LC firing rate may therefore differ in the context of different brain states associated with brain slice recordings, anesthesia, or wakefulness.

Indeed, LC firing rate is suppressed during slow wave sleep and REM sleep relative to the awake state (*37*). However, there is no reason to assume that LC firing rate differs between the anesthetized and awake states since sleep and anesthesia are not the same, mechanistically or phenomenologically (*38, 39*). Using the pooled dataset, we observed a main effect of experimental preparation (p<0.0001; **Figure 1O**; **Table S2**) in rats and mice (the two species for which we had both *ex vivo* and *in vivo* recordings). Pairwise comparisons showed that firing rates recorded under urethane were around 0.3 to 0.47 spikes/sec lower than those recorded under chloral hydrate (p<0.0001), isoflurane (p=0.006), and in slice (p=0.003) (**Figure 1O, Table S3**). Although firing rate differs between urethane and the other anesthetics, the firing rates of neurons in the awake state were similar to those recorded under anesthesia and in the deafferented brain slice. However, considering species as a factor modified the relationship between firing rate and brain state. The statistical model indicated an independent interaction between experimental preparation and species (p=0.001; **Figure 1P & Q, Table S2**). In rats, LC neurons from awake animals were more active than those recorded under urethane (p=0.034) or from brain slices (p=0.019). Surprisingly, mice showed a largely opposite pattern relative to what was observed in rats. Specifically, firing rate in awake mice was lower than LC activity under chloral hydrate (p<0.0001), isoflurane (p=0.0005), or urethane (p=0.001), or slice recordings (p<0.0001) (**Figure 1Q**; **Table S3**). The opposition between rats and mice is apparent in the effect size plots (**Figure 1P & Q**), which show positive effect sizes for rats and negative effect sizes for mice.

### The female LC has a suppressed firing rate in the awake state

Sex also modified the effect of brain state on LC firing rate. Our analysis revealed an independent interaction between experimental preparation and sex (p=0.004; **Figure 1R & S, Table S2**). In males, activity was similar across anesthetized, slice and awake animals except for small differences in firing rate between different types of anesthesia (**Figure 1R, see Table S3** for pairwise p-values). In contrast, firing rate was reduced in awake females compared to slice recordings and most anesthetics tested (**Figure 1S**; see **Table S3** for pairwise p-values). The effect sizes compared to the awake state varied from 0.96 to 2.49 spikes/sec. Thus, in awake females, LC firing rate is suppressed by a large magnitude (∼1 spikes/sec or greater) in comparison to LC neurons recorded in deafferenated brain slices or under anesthesia.

### Genetic modifications used in optogenetic and chemogenetic experiments may not strongly influence LC firing rates

In addition to the variety of experimental preparations used to study LC neuronal activity, genetic modifications enabling optogenetics and chemogenetics are used in neuroscience research. These genetic modifications typically involve expressing Cre recombinase throughout the brain (e.g., in all catecholaminergic neurons or in all GABA neurons). We compared recordings from *(i)* dopamine beta-hydroxylase (DBH)-Cre mice, which express Cre in noradrenaline-producing neurons (*14*); *(ii)* vesicular GABA transporter (VGAT)-Cre mice, which express Cre in GABAergic interneurons (*40*); and (iii) tyrosine hydroxylase (TH)-Cre rats, which express Cre in all catecholaminergic (dopaminergic and noradrenergic) neurons (*41*).

There was a significant effect of the type of genetic manipulation on firing rates (p<0.001, **Figure 1T, Table S2**). Post-hoc testing demonstrated that wild-type animals had higher firing rates compared to VGAT-Cre animals (p=0.0007) but were not different from DBH-Cre or TH-Cre animals (**Figure 1T**, **Table S3**). The effect of these genetic manipulations was not dependent on age or sex. Note that we could not assess any interaction effects with state or with species because VGAT-Cre and DBH-Cre data were from mice, while TH-Cre data were from rats (**Table S2**). This result may indicate that a genetic modification, which is used to enable optogenetic or chemogenetic manipulation of GABAergic interneurons throughout the brain, could reduce LC activity on the order of 0.51 spikes/sec compared to wild-type animals.

However, VGAT-Cre mice contributed a small number of neurons from a single laboratory. Further study of LC firing rate differences is necessary through the contribution of VGAT-Cre and wild-type mice and rats from a variety of laboratories.

### The LC becomes gradually more active over the course of aging

LC dysfunction is a universal feature of Alzheimer’s disease, Parkinson’s disease, and other neurodegenerative disorders beginning with the early aggregation of disease-specific proteins followed by late-stage degeneration (*19*). The largest risk for developing neurodegenerative disorders is age (*42*). Moreover, age heavily impacts LC structure and function, such as preclinical studies that have reported LC cell body and axon loss over the course of aging (*43–47*). We investigated whether there were any changes in LC firing rates during the process of normal aging. Rodents were designated as adolescent (<3 months), adult (3-12 months), or aged (>12 months), while all NHP data came from adult animals (10-16 years) (*48, 49*). There was a main effect of age on firing rates (p=0.006; **Figure 2A**, **Table S2**). Adolescent firing rates were lower than both adult (p=0.0002) and aged animals (p<0.0001), while LC activity in adults was lower than in aged animals (p=0.035) (**Figure 2A, Table S3**). Note that the interaction of age with species and state were not estimable (**Table S2**). There were no interactions between age and any other factor, suggesting that the gradual increase in LC activity during aging is not genotype- or sex-dependent. Thus, LC firing rates gradually increase with age, which has implications for understanding the contributions of LC activity to behavioral changes during aging and disease prevalence during aging.

**Fig. 2.**
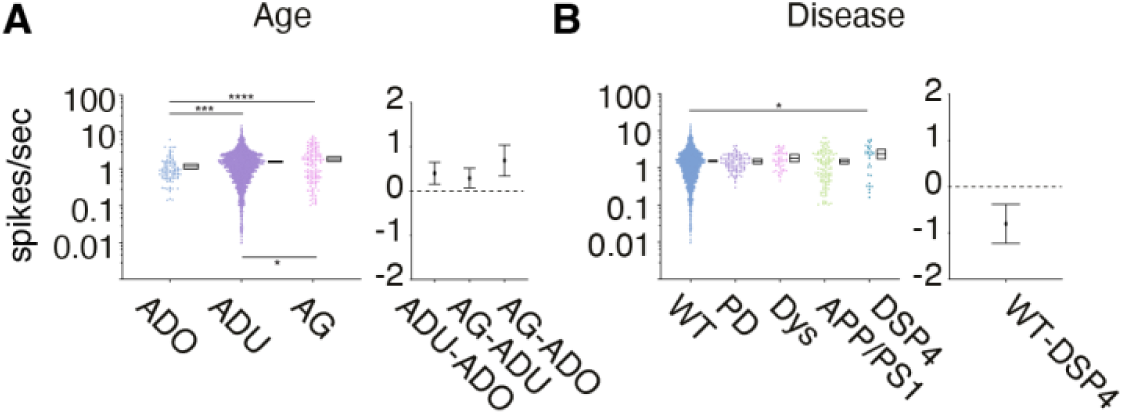
LC firing rate increases with age but does not change after genetic or pharmacological manipulations associated with neurodegenerative disease. All panels are formatted as in Fig. 1. Data points are individual neurons. All data are from rats and mice. (**A**) There was also a main effect of age on LC firing rates, with firing rate gradually increasing over the course of aging. (**B**) There was a main effect of disease model on LC firing rates and post-hoc analysis revealed that DSP-4 lesions increased LC activity. Abbreviations: ADO – adolescent, ADU – adult, AG – aged, WT – wild-type, PD – parkinsonian, Dys – dyskinetic. *p < 0.05, ***p < 0.001, ****p < 0.0001.

### LC firing rate is unaffected in models of neurodegenerative disease unless the degeneration specifically targets LC neurons

Given our finding that LC firing rate increases with age, which is a risk factor for developing neurodegenerative disorders, we wondered whether LC activity might be disrupted in genetic and pharmacological rodent models of neurodegenerative diseases. We analyzed the firing rate of neurons recorded from TgF344-AD rats that carry transgenes expressing autosomal dominant, early onset Alzheimer’s disease-causing APP/PS1 mutations (*50*). Importantly, these rats develop hyperphosphorylated tau in the LC prior to amyloid or tau pathology in forebrain regions (*43*), similar to the temporal pattern of tau deposition observed in humans (*51*). The dataset also included three pharmacological manipulations to induce neurodegenerative disease-like states. The first manipulation was the selective adrenergic neurotoxin, N-(2-chloroethyl)-N-ethyl-2-bromobenzylamine hydrochloride (DSP-4), which induces LC noradrenergic axonal degeneration (*52*). The second manipulation were rats unilaterally injected with 6-OHDA into the medial forebrain bundle to induce dopaminergic system neurodegeneration while sparing the noradrenergic system using pretreatment with desipramine (*53*). This second manipulation produces a hemiparkisonian disease model. The third manipulation combined 6-OHDA injections into the median forebrain bundle with 3 weeks of L-DOPA treatment, which produces a dyskinetic disease model (*54*). We asked whether these manipulations led to changes in LC firing rates that could be relevant for the presentation of neurodegenerative disorders. There was a main effect of disease model type on firing rates (p<0.0001; **Figure 2B, Table S2**). Post-hoc analysis revealed that animals treated with DSP-4 had higher firing rates than wild-type animals (p=0.0249; **Figure 2B, Table S3**). Unexpectedly, LC activity did not differ from control animals in any of the other disease models.

Due to the relationship between neurodegenerative disorders and age (*42*), the age of the animal might interact with the preclinical models to produce a change in LC activity. However, we found that the effect of these disease models on firing rate was not dependent on sex or age (**Table S2**). Note that interactions with species or brain state were not estimable. Overall, our findings are unexpected given that these disease models presumably relate to symptoms through altered noradrenergic neuromodulation by the LC but resulted in no change in baseline LC activity except for the DSP-4 pharmacological lesion model.

### LC activity spans a spectrum of diverse firing patterns that fall upon – but are constrained to – a low-dimensional manifold

We have shown that the time-averaged firing rates of LC neurons varies across species and multiple intrinsic and extrinsic biological factors. It is plausible that the patterns of LC activity may also vary. Traditionally, these patterns have been labeled as either a “tonic mode” or “phasic mode” of firing (*55–57*). Tonic spiking has been described as irregular discharging of spikes with inter-spike intervals (ISIs) lasting several hundreds of msec (resulting in a firing rate below ∼6 spikes/sec), while the phasic mode consists of 2-3 spikes within a 50-100 msec window (*57*) and correspondingly short (<80 ms) ISIs (*53, 58*). This two-mode dichotomy is used, not only in studies of single unit LC activity in animals, but also in human cognitive neuroscience studies where LC single neuronal activity is inaccessible, as well as in computational models of cognition and learning (*8–10*).

We detected the predominant time-averaged firing patterns across neurons by conducting a principal component analysis (PCA) of the logarithmically binned ISI distributions (ranging from 1 ms to 100 sec) across 1,708 neurons (see exclusion criteria from **Figure 1**). These 1,708 distributions characterized the time-averaged firing patterns from the entire recording of each neuron. The first three principal components (PC) explained 59.56% of the variance in the data with a drop to 7.89% at PC4 and <5% for all subsequent PCs. Thus, we plotted the first three PCs of the ISI distributions. We expected two discrete clusters of points that would correspond to neurons with irregular, infrequent spike times that either are, or are not, mixed with phasic bursts of spiking (< 80 ms ISIs). Instead, the firing patterns of individual neurons spread along a low-dimensional manifold with a dense central “core” area along the third PC zero axis and a “perimeter” area characterized by a dispersal of points in the positive and negative third PC axis (**Figure 3A** and **Supplementary Movie 1**). We sampled points from the core and perimeter and observed more rhythmic ISIs around perimeter area and irregular activity in the core (**Figure 3B-E, Supplementary Movie 2, Supplementary Movie 3**). In contrast to the notion that a single ISI cutoff divides LC activity into two firing patterns, we instead found that both rhythmic and irregular firing patterns spanned around 100 msec up to multiple seconds. This continuum of firing patterns among 1,708 LC neurons is evidence against tonic and phasic spiking modes that are characterized by rigid descriptors.

**Fig. 3.**
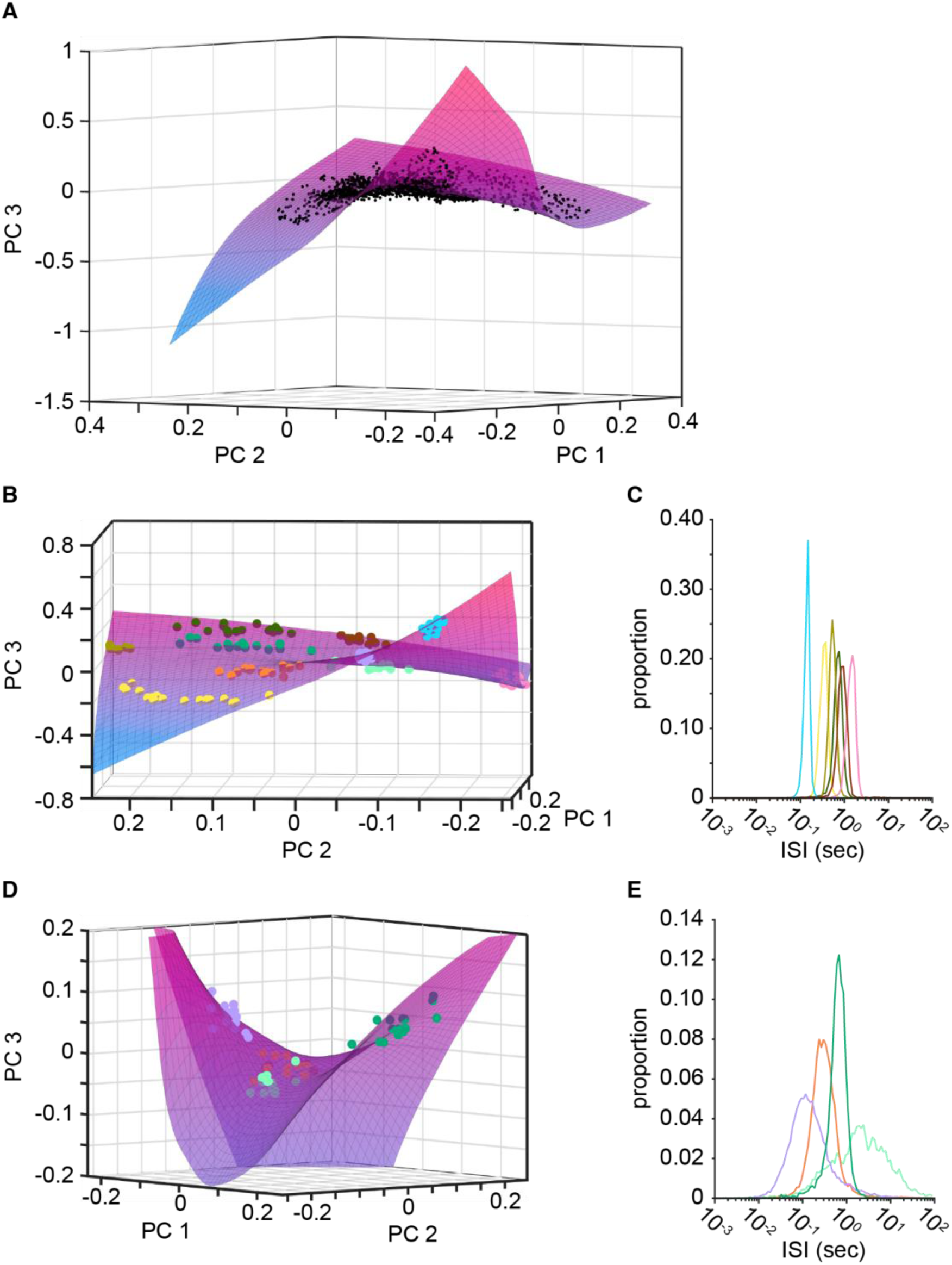
Neuronal firing patterns are distributed along a low-dimensional manifold characterized by diverse firing patterns. (**A**) The 3D scatter plot shows the principal components (PC) of each neuron on the PC1, PC2, and PC3 axis (N = 1,708 neurons). PCA was performed on the logarithmically-binned ISI distribution of each neuron. The pseudocolor surface shows the manifold fit to the points, which is colored according to the PC3 axis. The manifold consists of a dense, central core of neurons and additional neurons spanning the perimeter. (**B**) Subsets of 15 neurons were selected (at random) from different areas of the core and perimeter of the manifold. Neurons from different areas are plotted in different colors. (**C**) The average ISI distribution of the randomly selected neurons from the perimeter of the manifold (N = 15 neurons per average distribution). Note that the x-axis is logarithmically-scaled. (**D**) The PC3 axis of the manifold has been limited (±0.2) to focus on the central core region of the manifold. The different colored points are 15 randomly selected neurons in 4 different groups. Two groups (light green and orange) have negative PC3 values, but are separated in the PC1 and PC2 axes. Two groups (purple and dark green) have positive PC3 values and are separated in the PC1 and PC2 axes. These points are also all visible in the central area of the manifold in panel B. (**E**) The average ISI distributions for these central core groups (N = 15 neurons per average distribution). Note that the x-axis is logarithmically-scaled.

Specific LC activity patterns were not on the manifold, such as a highly positive PC2 and highly positive PC3. This suggests that some activity dynamics may not permissible (or not expressed in this dataset). Therefore, despite the diversity in the patterns of LC spiking, there are patterns which are not observed and may not occur naturally. Intriguingly, deafferanted brain slices, which made up one component of this dataset, are not natural. This begs the question of whether brain slice spiking patterns might map onto a specific area of the low-dimensional space. Therefore, we next assessed whether any factors (sex, species, age and brain state) were biased toward specific areas of the manifold.

### Firing patterns can occupy specific areas of the manifold when brain state, species, age, and genotype are considered

Brain slice firing patterns fell along the perimeter of the manifold, while firing patterns in awake organisms were concentrated in the core (**Figure 4A & B**). The firing patterns under anesthesia were spread across the entire manifold, but avoided the perimeter occupied by slice recordings. The difference in firing patterns between slice and awake recordings manifested as more rhythmic and with a lower limit of approximately 100 ms, while LC firing patterns during wakefulness exhibited comparably shorter (down to approximately 10 ms) as well as longer (up to approximately 10 sec) ISIs (**Figure 4C**). We used the two-sample Kolmogorov-Smirnov test to compare shapes of the ISI distributions. This test does not compare summary statistics (e.g., the mean ISI), rather it directly compares the distribution shapes by comparing the cumulative distribution functions. Awake recordings significantly differed from slice recordings (KS = 0.374, p<0.001), as did anesthetized recordings (KS = 0.353, p<0.001). Awake and anesthetized recordings had similar ISI distributions (KS = 0.111, p = 0.549). Overall, LC neurons recorded from brain slice exhibit clearly distinct firing patterns from awake and anesthetized recordings.

**Fig. 4.**
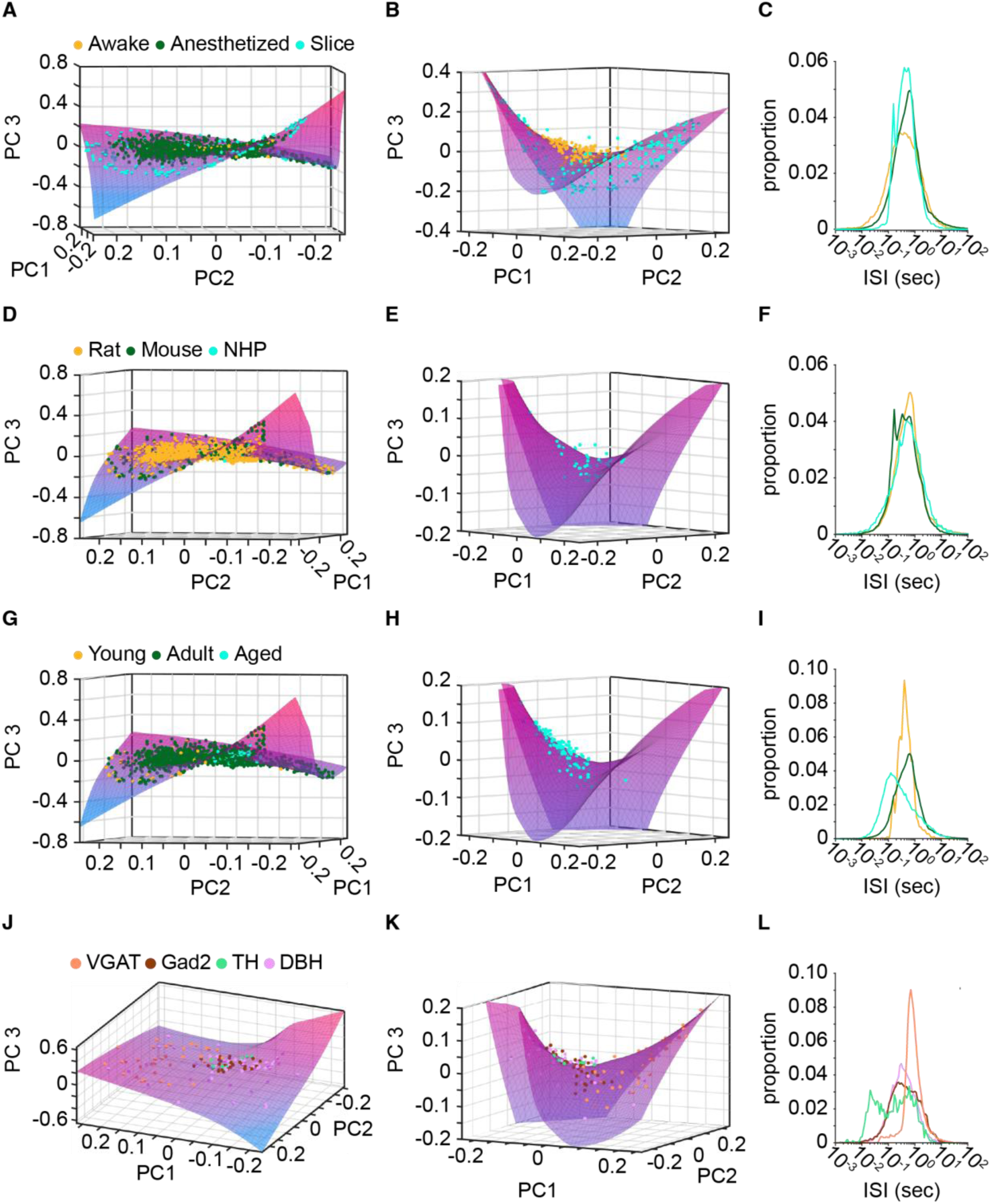
Intrinsic and extrinsic factors were biased toward particular areas of the manifold. (**A**) The PC scores of neurons recorded in the awake, anesthetized, or brain slice states are plotted on a 3D scatter plot of PC1, PC2, and PC3. N = 136 awake neurons; 1,376 anesthetized neurons; and 196 brain slice neurons. (**B**) The PC3 axis has been limited (±0.4) to show the central core area of the manifold. The neurons recorded in the awake state and brain slice state are shown. (**C**) The ISI distribution of all awake neurons (orange line, average of N = 136 neurons), all brain slice neurons (cyan line, average of N = 196 neurons), and all anesthetized neurons (green line, average of N = 1,376 neurons). Note that the x-axis is logarithmically-scaled. (**D - F**) The plots are the same format as panels A through C. The PC scores are shown for different species. N = 1,375 rat neurons; 292 mouse neurons; and 41 NHP neurons. In panel E), the PC3 axis of the manifold has been limited (±0.2) to show the central core area. The NHP neurons were constrained to this central area. Note that the x-axis is logarithmically-scaled. Differences between the distributions appear smaller on a log scale. (**G - I**) The plots are the same format as panels A through C. The PC scores are shown for different ages. N = 41 young animal neurons; 1,552 adult animal neurons; 115 aged animal neurons. In panel H), the PC3 axis of the manifold has been limited (±0.2) to show the central core area. The aged animal neurons were constrained to the negative PC1 axis and (mostly) positive PC3 axis of the central area. (**J - L**) The plots are the same format as panels A through C. The PC scores are shown for different genotypes. N = 57 DBH-Cre animal neurons; 32 Gad2-Cre animal neurons; 9 TH-Cre animal neurons; and 35 VGAT-Cre animal neurons. The VGAT-Cre animal neurons are lateralized on the manifold toward positive PC1 scores.

Species-specific and age-specific firing patterns were also observed. NHPs, mice, and rats all occupied large portions of the manifold (**Figure 4D**) with NHPs occupying the core of the manifold (**Figure 4E**). The ISI distributions for each species were thus similar (**Figure 4F**), although the firing patterns for mice and non-human primates may differ (KS = 0.202, p = 0.030). However, this may be due to NHPs being recorded only during wakefulness which we found was associated with the core area of the manifold (**Figure 4B**). The perimeter of the manifold contained both rat and mouse neurons, which corresponds with the contribution of slice recordings from both species. We also assessed whether rodent species might separate males and females on the manifold and found no evidence that sex was associated with different patterns of firing (**Figure S2**). While neurons recorded from adolescent and adult animals were spread across the manifold (**Figure 4G**), those recorded in aged animals were localized to one side of the core (**Figure 4H, Supplementary Movie 4**). Given that a large number of adult neurons were also concentrated in the core, it is unsuprising that the ISI histograms for adults and aged animals did not significantly differ (**Figure 4I**, KS = 0.131, p = 0.338). Although adolescent neurons appear to be isolated around the perimeter of the manifold and have a significantly different ISI distribution to both adult (KS = 0.394, p<0.001) and aged (KS = 0.343, p<0.001) animals, this is explained by adolescent neurons being recorded exclusively in slice recordings. *In vitro* recordings are typically done in brain slices from young animals. These results highlight the need for single laboratories to record LC single neuron activity in a variety of states (e.g., anesthetized and awake NHPs) and across a variety of ages (e.g., brain slices from young, adult, and aged animals). Despite the need for more data in some contexts, it is nevertheless apparent that aged neurons occupy a specific area of the manifold in this dataset.

Finally, we assessed how manipulation of genotype could affect where LC neuronal activity patterns fall upon this manifold. These genetic modifications expressed Cre recombinase throughout the brain with specificity for distinct neuron types. DBH-Cre targeted noradrenaline-producing neurons, TH-Cre targeted all catecholaminergic (dopaminergic and noradrenergic) neurons, and VGAT-Cre targeted GABAergic interneurons. VGAT-Cre neurons skewed toward one end of the manifold (**Figure 4J, Supplementary Movie 5**) and exhibited particularly rhythmic ISIs (**Figure 4K**). The ISI distribution for VGAT-Cre significantly differed from the distributions for TH-Cre (KS = 0.263, p = 0.002), DBH-Cre (KS = 0.323, p<0.001), and Gad2-Ires-Cre (KS = 0.242, p = 0.005). DBH-Cre also appeared to be more diffusely spread across the manifold relative to TH-Cre and the ISI distributions for those groups also significantly differed (KS = 0.273, p = 0.001). These results suggest that merely expressing Cre in a variety of neurons – brain wide – may alter the spontaneous firing patterns of LC neurons.

### The temporal dynamics of LC spike patterns exhibits a variety of attractor dynamics that are predominantly expressed according to species, age, sex, and genetic modification

Spike train patterns are dynamic over time. The PCA of ISI distributions (**Figures 3 and 4**) as well as the analysis of mean firing rate (**Figures 1 and 2**) were time-averaged analyses. We next characterized the second-order statistics of LC neuronal spiking using return maps (also known as Poincaré maps) which are visualizations of the joint relationship between successive inter-spike intervals. The return map can show temporal patterns including stable states (attractors) of neuronal firing (*59, 60*). **Figures 5A and 5B** illustrate this concept using synthetic data for two neurons, one which fires regularly and the other which fires tonically with occasional bursts. The return map distinguishes these two patterns and provides an intuitive visualization of the second-order statistics of spike trains. Here, we calculated return maps for the 1,708 LC neurons that met requisite conditions for ISI analysis. We observed several unique second-order spiking patterns, which we manually grouped. We then assessed how distinct types of second-order firing patterns mapped onto various factors.

**Fig. 5.**
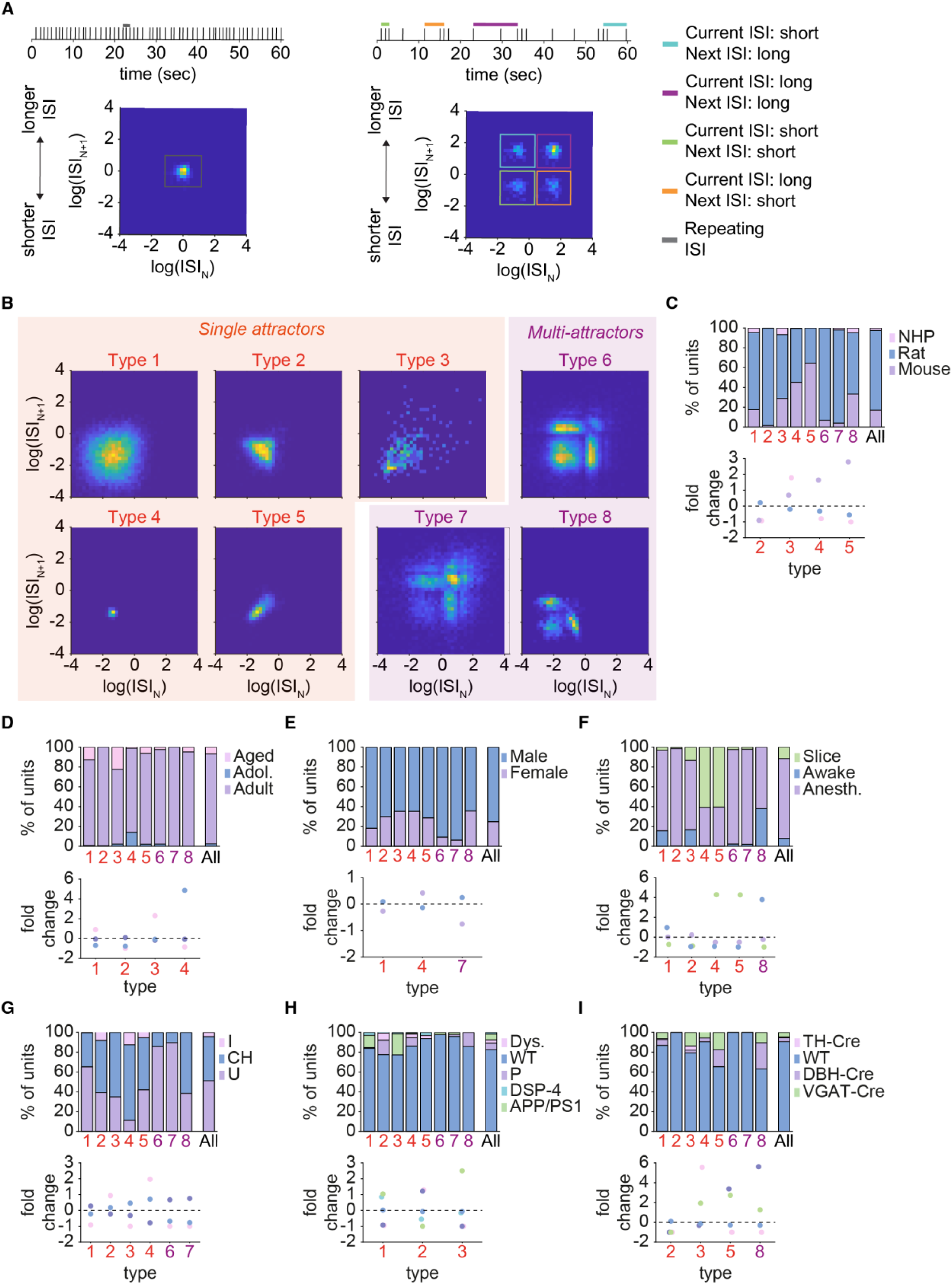
Intrinsic and extrinsic factors are associated with different types of attractor dynamics. The return density plots show the relationship between pairs of ISIs in a single neuron, namely the ISI of a given neuron on the x-axis against the following ISI of that same neuron on the y-axis. The frequency in which those relationships occur for a given unit are represented in the color where blue is no/infrequent ISI pairings and yellow is a pairing that occurs often. (**A-B**) Example neurons with their respective raster plot (top) and return density plots (bottom) show the relationship between firing rate, ISI, and return density plots. (**A**) Example neuron with a regular/tonic firing rate results in a return density plot with one point. (**B**) Example neuron with periods of tonic firing along with bursts results in a return density plot with a distinct shape of four points. Burst firing is highlighted in green in the raster and return density plot. Short ISIs followed by long ISIs (blue) and long ISIs followed by short ISIs (orange) are represented in the upper left and lower right corners of the return density plot. Tonic firing, long ISI followed by long ISIs, are highlighted in purple in the raster plot and return density plot. (**C-J**) Return density plot for the manually identified patterns fell into these 8 types. (**C-G**) Return density plots of Types 1 through 5 (**C:** N=689; **D:** N=560; **E:** N=90; **F:** N=206; **G:** N=48) show single attractor patterns, likely representing different forms of irregular tonic baseline activity. (**H-J**) Return density plots of Types 6 through 8 (**H:** N=44; **I:** N=50; **J:** n=21) contain multiple attractors, ranging in count from 3 to 5. Multi-attractor patterns have varying number of attractors, temporal gaps between attractors, and variability in the density around each attractor.

We identified 8 second-order firing patterns among the 1,708 neurons. **Figures 5C-J** shows an example unit from each of these 8 types of return maps. Types 1 through 5 (accounting for 62.5% of the patterns produced and most of the neurons) show a single stereotyped firing pattern (i.e., an attractor) with different shapes due to ISI variability **(Figures 5C-G)**. This pattern is expected from irregular tonic baseline activity. For example, the return density plots of type 1 and 4 are both evenly distributed around a common ISI value but the variability in type 1 is larger than type 4, which is highly discrete and the result of extremely regular activity similar to the synthetic raster used in **Figure 5A**. Types 6 through 8, on the other hand, have multiple attractors that also differ in their shape and variability **(Figures 5H-J)**. The neurons with multi-attractor firing patterns are generated by tonic firing interspersed with bursting at discrete relative frequencies. This identifies specific relative relationships in spontaneous bursting activity of LC neurons. The number of units in each type varied from 21 to 689. More complex multi-attractor firing patterns were observed in a minority of neurons (n=115, 6.73%), while most neurons fired in type 1 or type 2 single-attractor patterns (n=1249, 73.13%). Descriptive values for each type can be found in **Table S4**. These findings show that LC neurons produce firing patterns that are stereotyped attractors, and some neurons display transitions between multiple attractors.

Different extrinsic and intrinsic factors might bias LC neurons toward firing in particular second-order patterns. We tested this idea by characterizing to what degree different factors were expressed in each return map type by comparing the proportion of neurons with a specific factor to the overall population (**Table S4**). Our analysis revealed that some firing patterns are almost entirely expressed by a single species (**Figure 5K**). Type 2 (single attractor) was almost exclusively rat LC neurons. Types 4 and 5 (single attractor) had a greater proportion of neurons from mice and rarely included neurons recorded from NHPs. However, NHP LC neurons were overrepresented in type 3 (single attractor), which also contained a large proportion of mouse LC neurons. Therefore, we found species specific variations in tonic baseline firing intervals from mouse, rat, and NHP LC.

We also examined the effect of age on LC spike train dynamics (**Figure 5L**). Type 1 and 3, both single attractors with wide variability, had a higher proportion of aged animals. Adolescent neurons were especially common in type 4 which showed the most consistent ISIs. Type 2 was almost entirely made up of adult neurons with little/no adolescent or aged neurons. Thus, the adult LC may express a specific tonic firing pattern that distinguishes it from tonic activity patterns in adolescence or later life. Of note, variability (spread) between relative ISIs is very low in the type with more adolescent animals (**Figure 5F**; type 4), increasing in the adult animal type **(Figure 4D**; type 2), and the greatest variability is observed in aged animals **(Figure 5C & E**; types 1 and 3). Thus, distinct timepoints in development and aging are associated with specific LC second-order firing patterns. (Note, however, all LC neurons from adolescent animals were obtained from brain slice recordings. While aged animals had distinct firing patterns, the developmental transition of firing patterns from adolescence to adult animals may instead be due to the experimental preparation).

We investigated whether firing patterns were expressed to a greater extent in males compared to females and vice versa. However, no return map type was entirely specific to one of the sexes (**Figure 5M**). Male neurons predominated type 1 and 7, whereas type 4 had a higher proportion of female neurons. These results show that sex has some influence on second-order firing patterns, but all patterns are expressed in both sexes.

We next examined the role of experimental preparation on second-order dynamics (**Figure 5N**). Type 4 and 5 return maps (single attractors without much variability) had a significantly higher proportion of neurons recorded in a slice preparation and contained no neurons recorded in awake animals. Type 2 (single attractor) was almost entirely made up of neurons from anesthetized animals. Finally, type 1 (single attractor) and 8 (multiple attractors) were comprised of a higher proportion of neurons recorded in the awake state with a lower proportion of neurons recorded in the slice preparation. These results suggest that the preparation in which LC neurons are recorded is associated with specific second-order spike train statistics.

Given that experimental preparation is associated with altered LC spike train dynamics, we assessed whether the dynamics were influenced by the type of anesthetic agent utilized in anesthetized recordings (**Figure 4O**). Neurons recorded under urethane anesthesia were the majority in type 1, 6, and 7. Moreover, these types lacked neurons recorded under isoflurane. Similarly, there were no isoflurane neurons in type 3, but this type contained a higher proportion of chloral hydrate neurons. Finally, type 2 and 4 contained neurons recorded under isoflurane, as well as those recorded under chloral hydrate, in greater proportions. Thus, we found 75% of the second-order spike train patterns identified differed in their composition of neurons recorded under urethane, chloral hydrate, and/or isoflurane.

We tested whether models of neurodegenerative disease (i.e., genetic modification or pharmacological manipulations) were over-or under-expressed in any of the return map types. We found that type 2 had an increased proportion of neurons from pharmacological animal models of dyskinesia and Parkinson’s disease, but a decreased proportion of neurons from APP/PS1 genetic model and DSP-4 pharmacological model of neurodegenerative disease (**Figure 5P**). In contrast, we observed the opposite patterns in type 1 and 3. These clusters contained more APP/PS1 and DSP-4 neurons, but almost entirely lacked neurons from the pharmacological models of Parkinson’s disease and dyskinetic disorders.

Lastly, we assessed whether genetic manipulations for opto/chemo-genetics (Cre-expression) were differentially expressed across the 8 types of return density patterns (**Figure 5Q**). We found type 2 was almost entirely comprised of wild-type animals. Type 3 had a higher proportion of TH-Cre neurons, while type 5 and 8 had a greater proportion of neurons recorded DBH-Cre animals. Although not a large difference, types 3, 5, and 8 also had an increased proportion of neurons from VGAT-Cre animals. These results show that TH-Cre, DBH-Cre, and VGAT-Cre expressing neurons may each be biased toward specific relative firing patterns. Importantly, however, one type of firing pattern (type 2) was expressed only in wild-type subjects.

These multi-attractor types likely represent tonic and spontaneous phasic burst firing occurring in baseline LC activity. (**K-Q**) The bar plots (left panels) show the proportion of factors represented in each of the 8 return density types (type ID on x-axis) and the population as a whole (rightmost bar). The proportion of factors in each cluster is identified in each bar graph and was tested for differences compared to the overall population with a chi-squared test. Fold change in proportional representation (right panels) is shown for types where there was a significantly different proportion of factors. For each Type, the fold change was calculated by first subtracting the factor’s proportion in the population from the factor’s proportion within the Type. This difference was then divided by the proportion in the population. (**K**) Four types were comprised of neurons across species in different proportions compared to the population. Type 2 contained almost entirely neurons from rats. Mouse neurons were over-represented in Type 4 and Type 5.

NHP neurons were over-represented in Type 3. **L)** Age was associated with specific Types. Types 1 through 4 were comprised of significantly different proportions of neurons based on age. There was an increase in adolescent neurons in Type 4, adult neurons in Type 2, and aged neurons in Type 1 and Type 3. (**M**) Three of the Types (1,4, and 7) contained different proportions of neurons recorded from males or females. (**N**) Brain state varied greatly across the eight Types. Type 4 and Type 5 contained more neurons from slice recordings. Type 1 and Type 8 had more neurons from awake animals. Type 2 was almost entirely anesthetized neurons. (**O**) Almost all Types had different proportions of anesthetics compared to the overall population of neurons. (**P**) Neurodegenerative disease-associated manipulations were differentially represented in Type 1, Type 2, and Type 3. There was an over-representation of APP/PS1 and DSP-4 neurons in Type 1 and Type 3, and an over-representation of Parkinsonian and dyskinetic pharmacological manipulation neurons in Type 2. (**Q**) Type 3, Type 5, and Type 8 contained increased proportions of neurons from genetically manipulated (Cre) animals, while Type 2 was almost entirely wild-type animals. *p < 0.05, **p < 0.01 ***p < 0.001, ****p < 0.0001.

## Discussion

The LC is an ancestral brain region whose evolutionarily conserved function is thought to be achieved by two modes of LC firing: slow and irregular tonic activity or fast phasic burst activity. In contrast to this two-mode concept, we found that LC firing patterns spread along a low-dimensional manifold. The temporal dynamics of these firing patterns can be described as different types of attractors. Neither specific firing patterns, nor average activity levels, are conserved across species.

We used a negative binomial regression model to make a direct comparison of LC firing rate across species, and across other key intrinsic and extrinsic biological factors. The statistical model controlled for the individual and combined effects of various factors. Crucially, however, model-based significance testing was fully supported by assessments of effect size with 95% CIs. The 95% CI is an 83% prediction interval meaning it has an 83% chance of covering any future experiment’s mean effect size (*28, 30*). Thus, all findings were supported by both a statistical model and an approach not based upon p-values. We found that mean firing rate was higher in the mice compared to rats and NHPs. The model revealed that the effect of species upon mean firing rate also interacted with the brain state. The mouse LC was more active under anesthesia and in brain slices compared to the awake state, whereas in rats the opposite pattern was observed. Species and sex also interacted. In rats, neurons recorded from females were more active than those recorded in males, while the opposite pattern was observed in mice. It is known that the LC is a sexually dimorphic structure (*32*). Our findings may implicate the increased activity of female LC as a driver of sex-specific vulnerability to stress, anxiety, and substance-use disorder (*32, 61*), but our results also show that studies on the role of the LC in sex-specific vulnerabilities must be interpreted in a species-specific context. New data are needed to test whether species differences – specifically during wakefulness – could be due to sex of the animal, since neurons recorded during the awake state in mice came from females but, in rats and NHPs, they came from males.

It is important to consider that comparisons across factors inherently become comparisons across laboratories because the expertise of individual laboratories is usually centered upon one species and one experimental preparation (i.e., only awake, or with one type of anesthesia, or brain slices). While inter-laboratory variability in firing rate was low, we cannot exclude the possibility that some of the effects were due to differences in the procedures and techniques across laboratories. However, recent work comparing cortical, hippocampal, and thalamic single unit activity across 10 international laboratories found that average firing rate did not differ across laboratories if quality control criteria (similar to what we have employed in this study) are used (*62*). Nevertheless, resolving any inter-laboratory effect on our results will require individual laboratories to record diverse datasets by working across species, multiple experimental preparations, and recording from both sexes.

One of the most important observations from this large, pooled dataset is that LC firing patterns are highly diverse but still constrained onto a low-dimensional manifold. In other words, LC firing patterns spread along a spectrum of potential patterns, but some “off-manifold” patterns were not observed in this data set. The dynamics of these patterns can be described as different types of single attractors and multi-attractors. Studies of LC neuronal activity have defined tonic activity and phasic activity by dividing these firing modes using a threshold of either a firing rate >6 spikes/sec (*56, 57*), or ISI of less than 80 msec (*53, 58*). Our results suggest that, rather than using a rigid cut-off in firing rate or ISI to define two modes of activity, LC neuronal activity patterns must instead be appreciated as heterogeneous and falling across a spectrum. This has implications for human neuroimaging studies, cognitive psychology studies, and computational neuroscience studies which invoke the dual-mode conceptual framework (*8–10*). The broadening of LC modes of operation also has importance for understanding and interpreting pupillometry data as a correlate of LC activity (*63, 64*), which is commonly used to non-invasively index LC activity in human subjects. The heterogeneity of LC firing patterns also has implications for experiments probing LC function through optogenetic or electrical neuro-stimulation, which drive neurons at a specific timing and pattern (*65–68*). Stimulation parameters for these studies are consistently applied and do not reflect the underlying population heterogeneity reported here. Diversity in firing patterns is also important for understanding neuromodulatory functions of the LC throughout the central nervous system because when rodent LC neurons emit spikes at a short ISI, their axons can become depolarized so that subsequent spikes in the burst are conducted faster (*69*). Axonal acceleration of spikes may affect the magnitude of NE release (*70*) and the propensity for LC neurons to release co-transmitters (*15, 71, 72*).

We observed that firing patterns differed based on species, sex, age, and brain state. For instance, we observed firing patterns that were specific to rats and others that were specific to mice. Moreover, brain slice recordings were biased toward the perimeter of the manifold and the attractor dynamics were pacemaker-like (type 4 and type 5 Return plots), which may be explained by the severance of afferent connections and removed external sensory input relative to *in vivo* anesthetized and awake recordings . While pacemaker-like activity was most prominent in brain slices, such patterns were still observed in anesthetized and awake animals and are therefore not “unnatural” brain activity patterns. While the diversity in LC firing patterns is shared across different experimental preparations, some preparations are particularly appropriate for studying some areas of the manifold. The differential expression of diverse firing patterns across species and biological factors suggests that the neurotransmitter release properties and functional effects of LC neurons could have nuanced and previously unappreciated differences based on species, sex, age, and brain state.

Lastly, our findings have implications for understanding aging and neurodegenerative disorders. LC firing transitioned through different attractor dynamics from semi-rhythmic firing in the adolescent LC that becomes predominantly irregular in adulthood and finally more rhythmic, faster activity in aged animals. Mean firing rate gradually increased with age in multiple species, both sexes, and across *in vitro* recordings, *in vivo* anesthetized, and awake recordings. This is consistent with human neuroimaging (*73*) and mouse slice recordings (*74*) but discrepant with a prior anesthetized rat study which may be due to its smaller sample size (*75*). Age-related changes could be a compensatory adaptation to the NE fiber loss that occurs naturally with age in rodents and humans (*43, 44*).

LC cell body and fiber loss also occurs in neurodegenerative diseases (*19*). We compared multiple neurodegenerative disease-related genetic or pharmacological manipulations and found, among those models, a convergence upon single attractor spiking dynamics. The APP/PS1 and DSP-4 animals were overrepresented in type 1 and type 3 single attractor Return plots and underrepresented in type 2 single attractor Return plots, whereas dyskinetic and parkinsonian rat models had the opposite patterns. Our results suggest that single attractor (rather than multi-attractor) dynamics are a shared feature across neurodegenerative disorder-related LC modifications.

Using the shared contributions of data from multiple laboratories, our analysis uncovered diverse neuronal firing patterns, which can be described as a low-dimensional manifold. The dynamics of these firing patterns are expressed as attractors. Both firing patterns and average firing rates depended on intrinsic biological factors, such as species, sex, and age. Although the LC is an ancestral brain region shared by vertebrates, our data show that its activity has not been tightly conserved. The LC likely integrated into vertebrate nervous systems within the variable context of the evolutionary pressures that sculpted the nervous system of each species. Via its broad central nervous system-wide connectivity, this ancestral region may contribute to sex, species, and age-related differences in sensory processing, cognitive functions, motor control and behavior, as well as cardiovascular, immune, and metabolic function.

## Funding

This work was supported by funding afforded to the following people:

- Natural Sciences and Engineering Research Council of Canada PGS-D Fellowship (CJBM)
- National Institutes of Health grant DA10981 (CWB, DMD)
- National Institutes of Health grant MH14602 (CWB, DMD)
- Grant No. PID2022-142785OB-I00 funded by Ministerio de Ciencia (EB)
- Innovación y Universidades (MICIU)/Agencia Estatal de Investigacion (AEI)/10.13039/501100011033 (EB)
- ERDF A way of making Europe by the European Union Grant No. PDC2022-133987-100 funded by MICIU/AEI/10.13039/501100011033 (EB)
- European Union NextGenerationEU/PRTR Red Española de Investigación en Estrés/Spanish Network for Stress Research RED2022-134191-T financed by MICIU/AEI/10.13039/501100011033 (EB)
- CIBERSAM: CIBER -Consorcio Centro de Investigación Biomédica en Red CB07/09/0033 (EB)
- Young Investigator Award from Brain & Behavior Research Foundation (VBP)
- Future Leaders in Canadian Brain Research Program from Brain Canada (VBP)
- Natural Sciences and Engineering Research Council of Canada Discovery Grant RGPIN-2021-03284 (VBP)
- New Frontiers in Research Fund NFRFE-2022-00342 (VBP)
- Research Scholars Junior 1 Salary Award from Fonds de recherche du Québec Santé 311492 (VBP)
- National Institutes of Health grant R56-MH121918 (DJC)
- National Institutes of Health grant R21-DA058215 (DJC)
- Wellcome Trust PhD Programme in Neural Dynamics ref. 108899/Z/15 (OD, AS)
- National Institutes of Health grant T32-AA007573 (AD)
- National Institutes of Health grant F31-MH129112-01A1 (GD)
- National Alliance for Research on Schizophrenia and Depression Young Investigator Grant from the Brain and Behavior Research Foundation (RD)
- National Institutes of Health grant R01-DC012557 (RCF)
- National Institutes of Health Brain Initiative grant U19-NS107616 (RCF)
- NIDCD Predoctoral Fellowship F30-DC017351 (EG)
- National Institutes of Health grant R21-MH093904 (JIG)
- National Institutes of Health grant R01-NS101596 (XJ)
- National Institutes of Health grant R01-MH109556 (XJ)
- National Institutes of Health grant T32-EY07001 (XJ)
- KAKENHI grant 21H00219 (JPJ)
- Brain & Behavior Research Foundation NARSAD Young Investigator Award (APK)
- National Institutes of Health grant K08-MH122733 (APK)
- National Institutes of Health grant R21-MH134183 (APK)
- VA National Center for PTSD (APK)
- CT Department of Mental Health and Addiction Services (APK)
- National Institutes of Health grant F31-AG069502 (MAK)
- National Institutes of Health grant T32-MS96050 (MAK)
- National Institutes of Health grant R01-NS117899 (JGM)
- National Institutes of Health grant U01-AA020911-0851 (ZAM
- National Institutes of Health grant R01-DA049261 (ZAM)
- UNC Dissertation Completion Fellowship (ZAM)
- PID2 021-126434OB-I00 funded by MCIN/AEI/10. 13039/501100011033 (CM)
- ERDF A way of making Europe, Basque Government (IT1706-22; CM)
- Transborder Joint Laboratory (LTC) “non-motor Comorbidities in Parkinson’s Disease (CoMorPD)” (CM)
- Minister of Science and Technology Taiwan Grant Number: 105-2320-B-002-055-MY3 (MYM)
- CIHR Banting PDF BPF-151096 (MO)
- Wellcome Trust Senior Clinical Fellowship grant gr088373 (AEP)
- National Institutes of Health grant R01-DK098361 (AEP)
- National Institutes of Health grant R01-MH600670 (GRP)
- National Institutes of Health grant F31-MH131348 (ER)
- National Institutes of Health grant R01-MH119250 (SDS)
- National Health and Medical Research Council (GNT1193857; JMS)
- Australian Research Council (DP240101295; JMS)
- Bellberry Foundation (JMS)
- National Institutes of Health grant R01-MH126351 (MS)
- National Institutes of Health grant R01-MH133066 (MS)
- National Institutes of Health grant R01-MH085802 (MS)
- The Simons Foundation Autism Research Initiative through the Simons Center for the Social Brain (MS)
- Research Council of Finland PROFI6 funding program - UHBRAIN project (NKT)
- Start-up funding from the Helsinki Institute of Life Science (HiLIFE) at the University of Helsinki (NKT)
- National Institutes of Health BRAIN Initiative Grant 1R34NS123876 (NKT)
- JPMJFR2243 (AU)
- AMED-PRIME (AU)
- KAKENHI 21H05176 (AU)
- KAKENHI 22H02942 (AU)
- National Science Foundation GRFP grant DGE-2139839 (CRV)
- National Institutes of Health grant R01-MH112267 (QW)
- National Institutes of Health grant R01-NS119813 (QW)
- National Institutes of Health grant R21-MH125107 (QW)
- National Science Foundation grant IOS-2128543 (BDW)
- National Institutes of Health grant R01-MH101178 (BDW)
- National Institutes of Health grant R01-DA017960 (BDW)
- National Institutes of Health grant R01-NS32461 (BDW)
- National Institutes of Health grant U01-AA025481 (EV)
- National Institutes of Health grant R00-MH104716 (EV)
- National Institutes of Health grant R01-AG062581 (DW)
- National Institutes of Health grant RF1-AG079199 (DW)
- National Institutes of Health grant RF1-AG061175 (DW)
- Minister of Science and Technology Taiwan Grant Number 103-2302-B040-010-MY3 (HWY)

## Author contributions

Conceptualization: NKT

Data Curation: MAK, EM, LZ

Formal Analysis: MAK, BRM, ER, JMS, LZ

Investigation: VBP, CJBM, DJC, CCC, RFC, OD, DMD, AMD, GD, RD, ZF, EG, JCH, HI, MAK, JRK, CCK, YSK, RJL, YL, MLT, AMM, CAN, YRS, JR, ER, CR, ACS, RS, SDS, HCT, JAT, STS, AU, CRV, AV, HWY, JHY, HJY, LZ, ISZ

Methodology: MAK, ER, NKT, LZ

Project Administration: NKT, EV, DW

Resources: EA, EB, DMD, RCF, JIG, ZJ, JPJ, APK, JGM, ZAM, CM, MYM, AEP, SDS, MS, NKT, EV, QW, DW

Software: MAK, ER, NKT, LZ

Supervision: NKT, EV, DW

Validation: MAK, ER, NKT, LZ

Visualization: MAK, ER, NKT

Writing – Original Draft: MAK, ER, NKT

Writing – Review and Editing: EV, DW, GRP, BDW

## Competing interests

Qi Wang and Charles Rodenkirch are the founders of Sharper Sense, a company developing methods of enhancing sensory processing with neural interfaces. Alfred P. Kaye receives or has received research funding from Transcend Therapeutics and Freedom Biosciences, and has filed a provisional patent for combination psychedelic pharmacotherapies in PTSD. Anthony E Pickering reports research funding from Eli Lilly and is a member of the advisory board for Lateral Pharma. All other authors report no disclosures or conflicts of interest.

## Data and materials availability

All data are available on a public repository (DANDI Archive) [link will be added upon being sent for peer review]. The code for analysis is available on github [link will be added upon being sent for peer review].

## Materials and Methods

### Ethical guidelines

All animal studies in the United States were performed with approval from the Institution Animal Care and Use Committee at Baylor College of Medicine, Cold Spring Harbor, Columbia University, Emory University, Massachusetts Institute of Technology, New York University, Rowan University, University of North Carolina Chapel Hill, University of Pennsylvania, University of Wisconsin-Madison, Washington University School of Medicine, University, and Yale University in accordance with the National Institutes of Health Guide for the Care and Use of Laboratory Animals. Approval for animal studies outside of the United States were approved by the Animal Care and Experimentation Committee of the Institute for Research in Fundamental Sciences, the Animal Care and Use Committees of the RIKEN Brain Science Institute, the Institutional Animal Care and Use Committee of National Taiwan University, the Institutional Animal Care and Use Committee of Chung-Shan Medical University, the University of Bristol Animal Welfare and Ethical review body according to the UK Animals (Scientific Procedures) Act 1986 (license PPL3003362), the Committee for Animal Experimentation at the University of Cadiz and University of the Basque Country (UPV/EHU), in compliance with the European Community Guidelines for the Care and Use of Laboratory Animals (2010/63/EU) and Spanish Law for the protection of animals used for research experimentation and other scientific purposes (RD 53/2013), the Committee for Animal Experimentation at the University of Cadiz, The Regierungspräsidium Tuebingen & The Regional State Administrative Agency for Southern Finland according to German Law for the Protection of Animals in experimental research (Tierschutzversuchstierverordnung) & the European Community Guidelines for the Care and Use of Laboratory Animals (EU Directive 2010/63/EU).

### Data Collection and Organization

Data was sourced from various labs and imported into a custom structure in Matlab (MathWorks; version R2019a) for analysis. We acquired data on species, sex, strain, state (awake, slice, anesthesia and type), age, pharmacology and events (type and timestamp), and genotype. LC recordings were imported as timestamps of each individual spike within a given recording. Firing rates were calculated as the number of spikes divided by the timestamp of the last recorded spike.

### Statistical analysis of firing rates

Descriptive statistics were used to describe the characteristics of the sample of firing rates. Given the multitude of variables present in our analyses, we fit a regression to explain the individual and combined effects of factors on firing rates. Initial examination of the data revealed overdispersion. We therefore chose a negative binomial regression model, which is commonly used to model overdispersed data. Negative binomial regression is a generalization of a Poisson regression with less restrictive assumptions of equal mean and variance which was also present in our data. Given that we wanted to leverage cross-lab comparisons, we removed lab as a random effect and fit a marginal model instead. This was further justified by the large size of our dataset and model fit statistics (ratio of the deviance to degrees of freedom and dispersion parameter). A generalized estimating equation algorithm was then used to estimate the independent effects of individual factors and their interactions on firing rates (SAS PROC GENMOD).

Generalized estimating equations (GEE) are one popular algorithm for fitting marginal models when outcome is a discrete variable. Using GEE with offset of measurement time, the independent effect of age, sex, species, genotype and anesthesia/state as well as their effect modification on firing rate were assessed. If a significant effect modification (interaction) was present, estimated firing rate was compared at a different level of the components of the interaction. If the interaction was not significant, estimated firing rate was reported with adjustment of other features. GEE was also applied to the comparison of firing rate between single or various combination of methods which were used to verify LC neurons. To correct for multiple comparisons, a false-discovery rate of < 20% was applied and Benjamini-Hochberg procedure was used. All analyses were performed in SAS® 9.4 (SAS Institute Inc). All tests performed were two-tailed. Effect size plots showing the difference between the estimated means ± 95% confidence interval of the estimate was generated using the DABEST toolbox in MATLAB.

### Principal components analysis of inter-spike interval (ISI) distributions

ISIs were calculated by subtracting the time of each spike from the following spike time (spikex+1-spikex). The calculation was performed from the first spike recorded until the last spike of the recording prior to a pharmacological manipulation or external event (e.g., a sensory stimulus), if either was present. ISI histograms (calculated in 100 logarithmically-spaced bins from 1 msec to 100 sec) were analyzed using PCA. A smooth surface (i.e., manifold) was fit to the first three PCs using the MATLAB (v2025a) function, loess. The root mean square deviation was smaller for loess compared to the lowess function.

### Return maps

Second-order ISI patterns were visualized using return maps. These density plots show the ISI of a single neuron on the x-axis plotted against the following ISI of that same neuron on the y-axis. A two-dimensional histogram was created for each unit (ISIn x ISIn+1) binned in 40 bins on each axis centered around the maximum value in the two-dimensional histogram (±5) plotted on a logarithmic scale for better visualization. The bin with maximum density was centered in the plot to visualize the shape of the return map. We manually clustered the patterns into 8 types. We compared the proportion of a given factor within each return map type to the overall population of neurons in the dataset. Differences in the frequency of a given factor in each return map type were compared to the frequency of the factors in the total population of neurons. The proportion of neurons was compared using the chi-squared test with Bonferroni correction for multiple comparisons.

### Quality control of single unit recordings across labs

A total of 2,944 neurons were submitted across 20 labs. We discarded neurons for which more than 2% of the interspike intervals were less than 10 msec as described in prior work (76). This excluded 319 neurons from further analysis. Next, we removed spiking occurring during pharmacological manipulation with the exception of anesthesia. We sought to characterize only temporally stable recordings by removing neurons which did not spike more than 30 times throughout the recording (457 neurons were removed). At the request of the contributors of those data we also removed 199 neurons that were recorded from animals subjected to a stressor. Additionally, we removed 113 neurons in which the sex of the animal was unknown; this was necessary because the negative binomial regression model required each neuron to have information about each factor that could affect firing rate. These exclusions left only one neuron recorded under pentobarbital anesthesia, which was removed from subsequent analyses. After submitting data to this quality control procedure, a total of 1,855 neurons remained for analysis of firing rates. The number of neurons for each factor are reported in **Table S1**. Finally, since some recordings were provided with stimulus-evoked spiking following a stimulus-free baseline epoch, we determined for each neuron whether there was significant difference in the mean firing rate between the epoch of the recording without stimulation and the subsequent recording epoch containing sensory stimuli.

We reasoned that, since phasic LC responses to stimuli are transient biphasic events lasting ∼1000 msec, such transient changes would not lead to an overall change in firing rate averaged over a comparably much larger time window (the duration of this subset of recordings ranged from ∼5 seconds to ∼7 hours, and the median duration was 444 sec). We performed a Student’s t-test comparing multiple samples of firing rate (split into 10 equal time bins) between the recording epochs with, or without, sensory stimuli. If the introduction of sensory stimuli led to a significant (p<0.05) alteration of time-averaged firing rate compared to the preceding stimulus-free epoch, then the spike train of that neuron was analyzed only until the first stimulus and the remaining data were discarded. Out of 1,855 neurons, 560 were exposed to sensory stimuli; of those 560, we discarded the recording epoch with stimuli for 352 neurons. In addition to analyzing firing rate, we also analyzed firing patterns and, for that analysis, only included spike trains of individual neurons prior to any stimuli. Further, we excluded units that did not have at least 100 interspike intervals (ISIs) prior to any stimulus or pharmacology application but included neurons from animals with an unreported sex to increase sample size (1,708 neurons analyzed).

### Factors excluded from the model of firing rates

Animal strain was not included in the model because strain could be completely encompassed by another factor, in this instance species (i.e., Fischer animals are only in rats, C57Bl/6 animals are only mice). For similar reasons, the interactions between age and species (adolescent and aged neurons were only obtained from rats), age and anesthesia (aged animals were all recorded under chloral hydrate anesthesia while all adolescent animals were recorded in slice), species and genotype (specific genotypes came from the same species, such as APP/PS1 animals all being rats), and genotype and anesthesia (neurons from specific genotypes were recorded under the same anesthetic) could not be estimated.

**Fig. S1.**
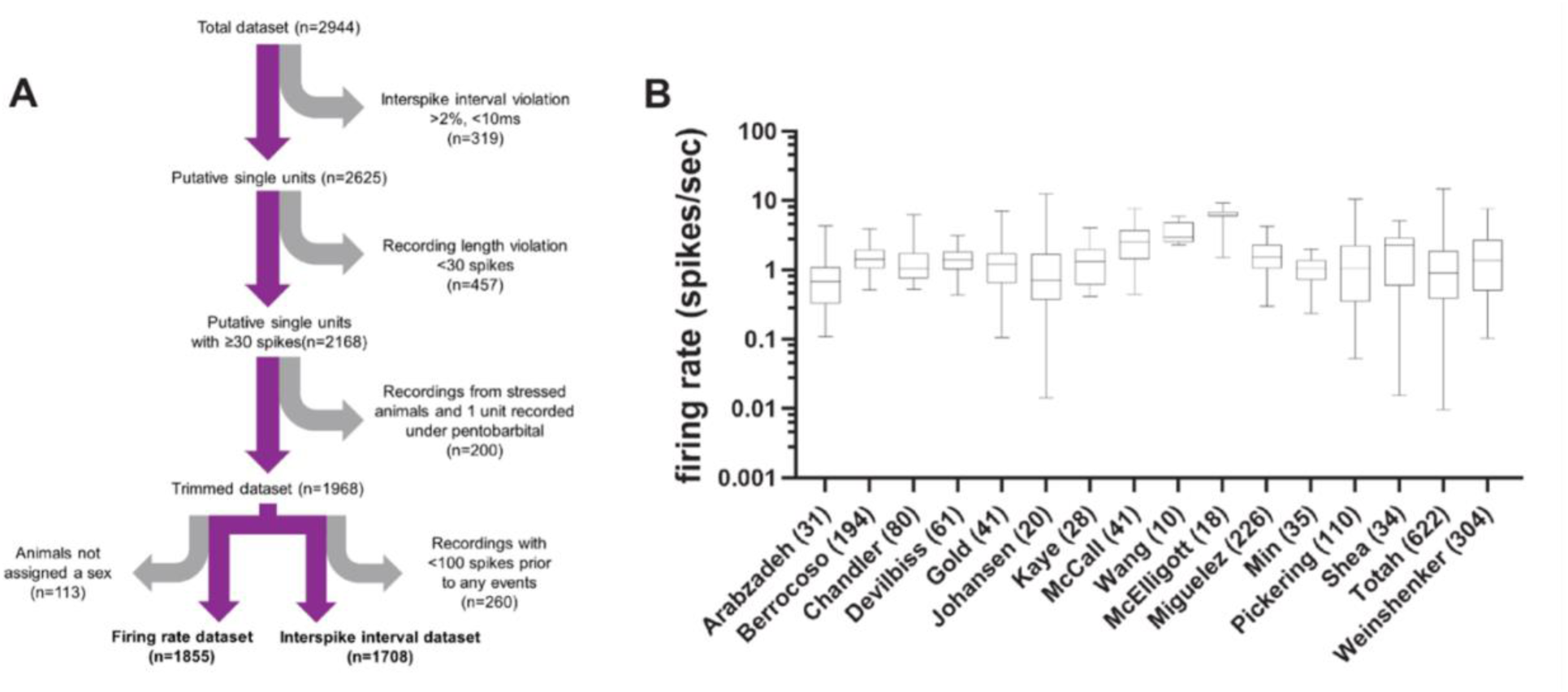
Quality control of the dataset and quantification of across lab variability in firing rate. (**A**) A flowchart visualizing the quality control steps of submitted data. Grey arrows indicate removal of neurons from the data set and the number of neurons removed is in parentheses. The number reported below each purple arrow shows the number of remaining neurons subjected to the next step in the quality control process. To maintain the largest appropriate dataset, slightly different criteria were used to qualify neurons be included for firing rate analysis (left, N=1,855) vs. interspike interval analysis (right, N=1,708). (**B**) The distributions of log-transformed firing rates of 1,855 neurons assessed in the firing rate analysis are shown separately by lab for the purpose of visualizing firing rate variability within and across labs. The average firing rate of each neuron contributed a data point. The whiskers plots show the minimum and the maximum, while the boxes indicate the 25%, 50%, and 75% bounds of the distribution. The x-axis lists the lab name and the number of neurons analyzed from that lab.

**Fig. S2.**
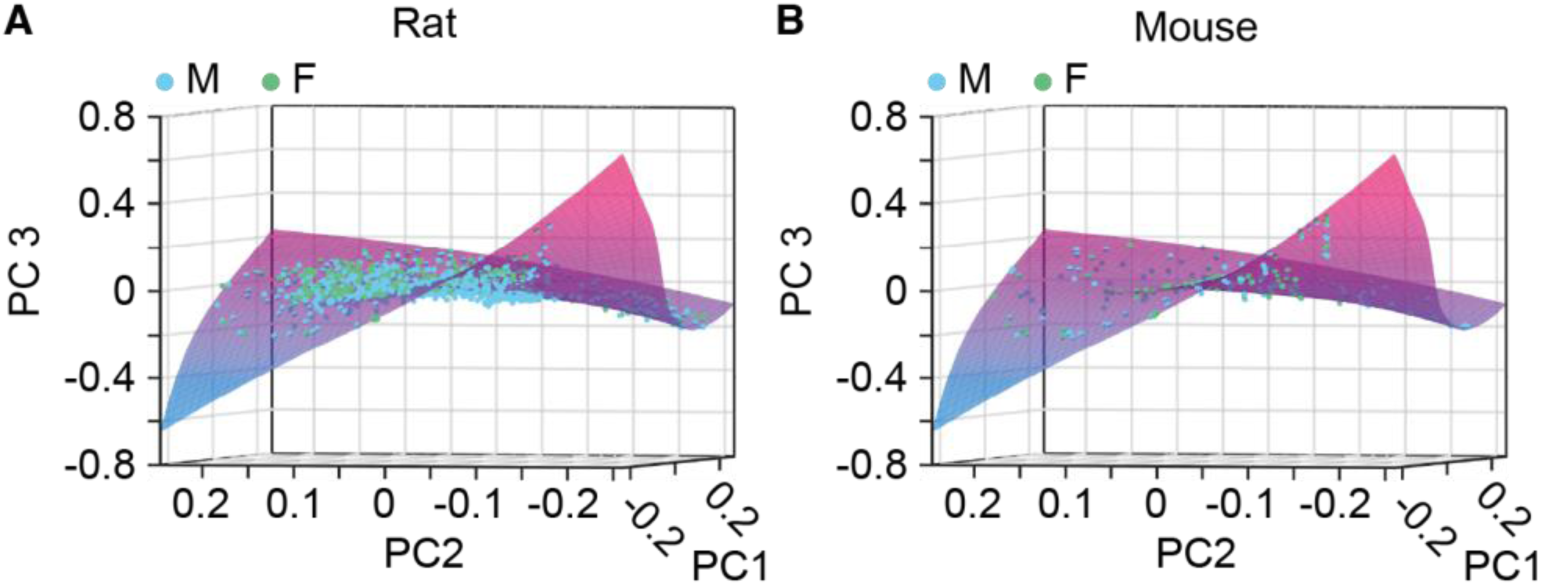
The distribution of male and female neurons on the manifold for mice and rats. The PC scores of neurons recorded in the awake, anesthetized, or brain slice states are plotted on a 3D scatter plot of PC1, PC2, and PC3. (**A**) Male and female rats. (**B**) Male and female mice.

**(see excel file for Table S1)**

**Table. S1.** Summary of the characteristics of the dataset after quality control. The percentages are based upon the total number of neurons subjected to analysis of firing rate (n = 1,855). WT refers to wild-type animals that were unmanipulated and not on another genetic background (i.e. cre-lines). Rodents were designated as adolescent (<3 months), adult (3-12 months), or aged (>12 months), while all NHP data came from adult animals (10-16 years) (46, 47).

**Table S2.**
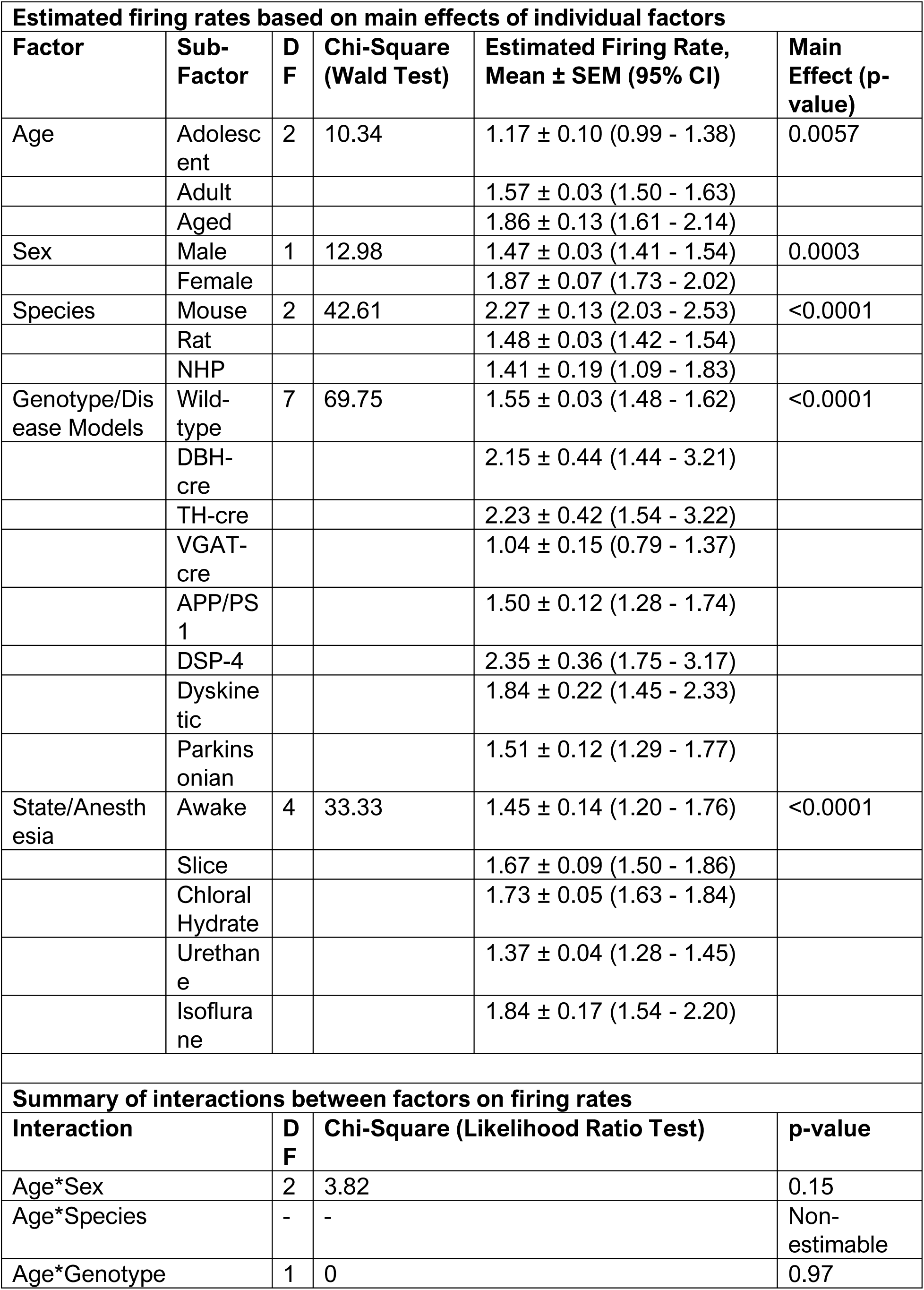
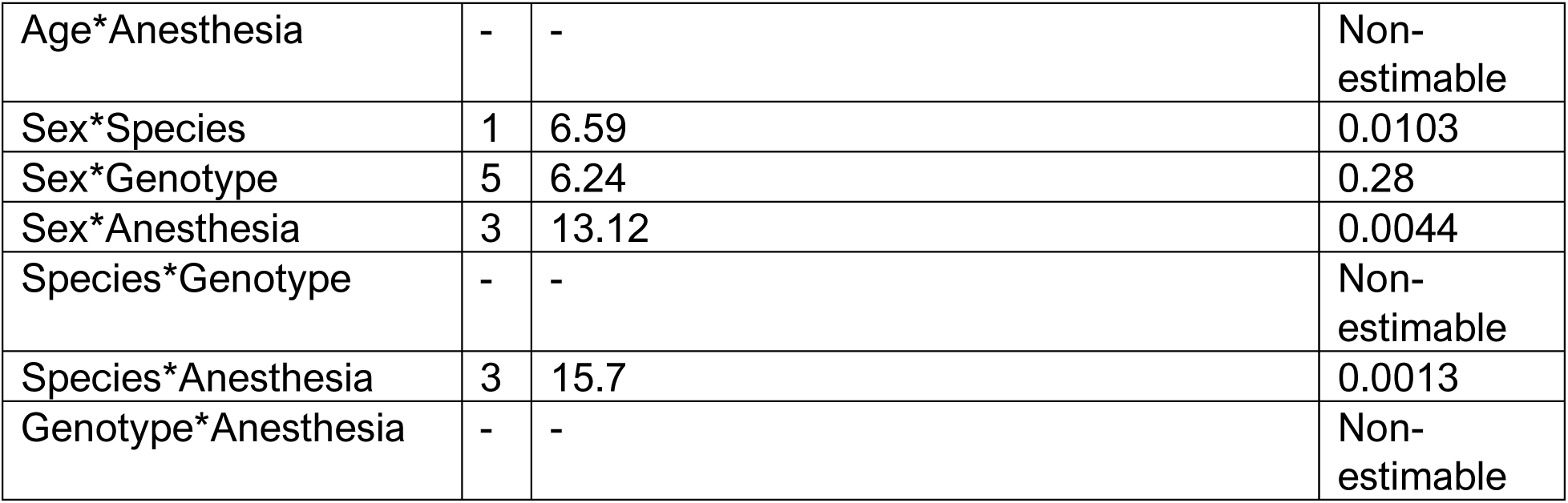
Detailed results of the negative binomial regression model of firing rates.

**Table S3.** Detailed results the post-hoc pairwise comparisons of differences in firing rates.

| Post-hoc pairwise comparisons of firing rates based on main effect of species |  |  |  |  |  |  |
| --- | --- | --- | --- | --- | --- | --- |
| Comparison | DF | Chi-Square (Wald Test) | Effect Size (95% CI) | p-value |  |  |
| Rat vs. Mouse | 1 | 36.537 | -0.79 (-0.97 – -0.61) | <0.0001 |  |  |
| NHP vs. Mouse | 1 | 14.45 | -0.86 (-1.49 – -0.23) | 0.0001 |  |  |
| Rat vs. NHP | 1 | 0.135 | 0.07 (-0.44 – 0.30) | 0.71 |  |  |
| Post-hoc pairwise comparisons of firing rates based on the interaction between species and experimental preparation |  |  |  |  |  |  |
|  | Effect Size (95% CI) |  |  | p-value (DF, Chi-Square [Wald Test]) |  |  |
| Experimental Preparation | Rat vs Mouse | NHP vs Mouse | Rat vs NHP | Rat vs Mouse | NHP vs Mouse | Rat vs NHP |
| Awake | 1.34 (0.30 – 2.38) | 0.52 (-0.15 – 1.19) | 0.82 (0.02 – 1.61) | 0.0036 (1, 8.472) | 0.0491 (1, 3.871) | 0.08 (1, 3.167) |
| Chloral Hydrate | -0.56 (-0.91 – -0.21) | - | - | 0.0206 (1, 5.361) | - | - |
| Isoflurane | -1.47 (-2.33 – -0.61) | - | - | 0.0170 (1, 5.697) | - | - |
| Slice | -1.00 (-1.49 – -0.60) | - | - | <0.0001 (1, 26.456) | - | - |
| Urethane | -2.57 (-3.20 – -1.94) | - | - | 0.0049 (1, 7.913) | - | - |
| Post-hoc pairwise comparisons of firing rates based on the interaction between species and sex (males) |  |  |  |  |  |  |

| Comparison | DF | Chi-Square (Wald Test) | Effect Size (95% CI) | p-value |
| --- | --- | --- | --- | --- |
| Rat vs. Mouse | 1 | 35.57 | -1.09 (-1.30 – -0.88) | <0.0001 |
| NHP vs. Mouse | 1 | 17.1 | -1.07 (-1.78 – -0.36) | <0.0001 |
| Rat vs. NHP | 1 | 0.02 | -0.02 (-0.35 – 0.31) | 0.89 |
| **Post-hoc pairwise comparisons of firing rates based on the interaction between species and sex (females)** |  |  |  |  |
| Comparison | DF | Chi-Square (Wald Test) | Effect Size (95% CI) | p-value |
| Rat vs. Mouse | 1 | 0.42 | -0.12 (-0.47 – 0.23) | 0.52 |
| **Post-hoc pairwise comparisons of firing rates based on main effect of sex** |  |  |  |  |
| Comparison | DF | Chi-Square (Wald Test) | Effect Size (95% CI) | p-value |
| Male vs. Female | 1 | 24.61 | -0.40 (-0.53 – -0.27) | <0.0001 |
| **Post-hoc pairwise comparisons of firing rates based on the interaction between sex and experimental preparation (male vs. female)** |  |  |  |  |
| Comparison | DF | Chi-Square (Wald Test) | Effect Size (95% CI) | p-value |
| Awake | 1 | 8.309 | 0.75 (0.00 – 1.50) | 0.0039 |
| Chloral Hydrate | 1 | 4.617 | -0.24 (-0.45 – -0.03) | 0.0317 |
| Isoflurane | 1 | 1.988 | -1.65 (-2.96 – -0.34) | 0.16 |
| Slice | 1 | 5.105 | -0.48 (-0.86 – -0.09) | 0.0239 |
| Urethane | 1 | 2.015 | -3.1 (-4.35 – -1.85) | 0.16 |

| **Post-hoc pairwise comparisons of firing rates based on the interaction between sex and species (rats)** |  |  |  |  |
| --- | --- | --- | --- | --- |
| **Comparison** | **DF** | **Chi-Square (Wald Test)** | **Effect Size (95% CI)** | **p-value** |
| Male vs. Female | 1 | 26.76 | -0.45 (-0.59 – -0.31) | <0.0001 |
| **Post-hoc pairwise comparisons of firing rates based on the interaction between sex and species (mice)** |  |  |  |  |
| **Comparison** | **DF** | **Chi-Square (Wald Test)** | **Effect Size (95% CI)** | **p-value** |
| Male vs. Female | 1 | 4.49 | 0.52 (0.02 – 1.02) | 0.034 |
| **Post-hoc pairwise comparisons of firing rates based on main effect of experimental preparation** |  |  |  |  |
| **Comparison** | **DF** | **Chi-Square (Wald Test)** | **Effect Size (95% CI)** | **p-value** |
| Awake vs. Chloral Hydrate | 1 | 3.376 | -0.28 (-0.60 – 0.04) | 0.07 |
| Awake vs. Isoflurane | 1 | 3.116 | -0.39 (-0.83 – 0.05) | 0.08 |
| Awake vs. Slice | 1 | 1.705 | -0.22 (-0.57 – 0.13) | 0.19 |
| Awake vs. Urethane | 1 | 0.352 | 0.08 (-0.18 – 0.34) | 0.55 |
| Chloral Hydrate vs. Isoflurane | 1 | 0.385 | -0.11 (-0.42 – 0.20) | 0.53 |
| Chloral Hydrate vs. Slice | 1 | 0.281 | 0.06 (-0.14 – 0.26) | 0.6 |
| Chloral Hydrate vs. Urethane | 1 | 27.743 | 0.36 (0.23 – 0.49) | <0.0001 |

| Isoflurane vs. Slice | 1 | 0.756 | 0.17 (-0.24 – 0.47) | 0.38 |
| --- | --- | --- | --- | --- |
| Isoflurane vs. Urethane | 1 | 7.482 | 0.47 (0.21 - 0.73) | 0.0062 |
| Slice vs. Urethane | 1 | 9.037 | 0.30 (0.13 – 0.47) | 0.0026 |
| **Post-hoc pairwise comparisons of firing rates based on the interaction between experimental preparation and sex (males)** |  |  |  |  |
| Comparison | DF | Chi-Square (Wald Test) | Effect Size (95% CI) | p-value |
| Awake vs. Chloral Hydrate | 1 | 0.02 | 0.03 (0.33 – 0.39) | 0.89 |
| Awake vs. Isoflurane | 1 | 0.13 | -0.09 (-0.57 – 0.30) | 0.71 |
| Awake vs. Slice | 1 | 0.37 | 0.12 (-0.26 – 0.51) | 0.54 |
| Awake vs. Urethane | 1 | 2.47 | 0.29 (0 – 0.58) | 0.12 |
| Chloral Hydrate vs. Isoflurane | 1 | 0.41 | -0.12 (-0.45 – 0.21) | 0.52 |
| Chloral Hydrate vs. Slice | 1 | 0.61 | 0.09 (-0.16 – 0.34) | 0.43 |
| Chloral Hydrate vs. Urethane | 1 | 9.65 | 0.26 (0.11 – 0.41) | 0.0019 |
| Isoflurane vs. Slice | 1 | 1.21 | 0.21 (-0.15 – 0.57) | 0.27 |
| Isoflurane vs. Urethane | 1 | 4.92 | 0.37 (0.12 – 0.64) | 0.0266 |
| Slice vs. Urethane | 1 | 2.084 | 0.17 (-0.02 – 0.36) | 0.15 |

| **Post-hoc pairwise comparisons of firing rates based on the interaction between experimental preparation and sex (females)** |  |  |  |  |
| --- | --- | --- | --- | --- |
| **Comparison** | **DF** | **Chi-Square (Wald Test)** | **Effect Size (95% CI)** | **p-value** |
| Awake vs. Chloral Hydrate | 1 | 21.11 | -0.96 (-1.74 – -0.18) | <0.0001 |
| Awake vs. Isoflurane | 1 | 4.49 | -2.49 (-4.10 – -0.88) | 0.0341 |
| Awake vs. Slice | 1 | 17.08 | -1.11 (-2.00 – -0.22) | <0.0001 |
| Awake vs. Urethane | 1 | 2.65 | -3.56 (-5.57 – -1.55) | 0.10 |
| Chloral Hydrate vs. Isoflurane | 1 | 1.73 | -1.53 (-2.79 – -0.27) | 0.19 |
| Chloral Hydrate vs. Slice | 1 | 0.57 | -0.16 (-1.38 – 1.08) | 0.45 |
| Chloral Hydrate vs. Urethane | 1 | 1.42 | -2.60 (-4.37 – -0.83) | 0.23 |
| Isoflurane vs. Slice | 1 | 1.37 | 1.38 (-0.12 – 2.88) | 0.24 |
| Isoflurane vs. Urethane | 1 | 0.19 | -1.07 (-6.32 – 4.18) | 0.66 |
| Slice vs. Urethane | 1 | 1.25 | -2.45 (-4.53 – -0.37) | 0.26 |
| **Post-hoc pairwise comparisons of firing rates based on the interaction between experimental preparation and species (rats)** |  |  |  |  |
| **Comparison** | **DF** | **Chi-Square (Wald Test)** | **Effect Size (95% CI)** | **p-value** |
| Awake vs Chloral Hydrate | 1 | 1.679 | 0.55 (-0.03 – 1.13) | 0.2 |
| Awake vs Isoflurane | 1 | 2.03 | 0.64 (-0.10 – 1.38) | 0.15 |
| Awake vs Slice | 1 | 5.523 | 1.01 (0.43 – 1.59) | 0.0188 |
| Awake vs Urethane | 1 | 4.486 | 0.89 (0.40 – 1.38) | 0.0342 |
| Chloral Hydrate vs Isoflurane | 1 | 0.291 | 0.09 (-0.23 – 0.41) | 0.59 |
| Chloral Hydrate vs Slice | 1 | 17.046 | 0.46 (0.21 – 0.71) | <0.0001 |
| Chloral Hydrate vs Urethane | 1 | 24.686 | 0.34 (0.22 – 0.46) | <0.0001 |
| Isoflurane vs Slice | 1 | 3.778 | 0.37 (0.02 – 0.72) | 0.05 |
| Isoflurane vs Urethane | 1 | 2.209 | 0.25 (-0.2 – 0.52) | 0.14 |
| Slice vs Urethane | 1 | 1.258 | -0.12 (-0.33 – 0.09) | 0.26 |
| **Post-hoc pairwise comparisons of firing rates based on the interaction between experimental preparation and species (rats)** |  |  |  |  |
| **Comparison** | **DF** | **Chi-Square (Wald Test)** | **Effect Size (95% CI)** | **p-value** |
| Awake vs. Chloral Hydrate | 1 | 19.763 | -1.35 (-2.36 – -0.34) | <0.0001 |
| Awake vs. Isoflurane | 1 | 12.074 | -2.17 (-3.57 – -0.77) | 0.0005 |
| Awake vs. Slice | 1 | 27.122 | -1.33 (-2.30 – -0.36) | <0.0001 |
| Awake vs. Urethane | 1 | 10.478 | -3.02 (-4.81 – -1.23) | 0.0012 |
| Chloral Hydrate vs. Isoflurane | 1 | 1.637 | -0.82 (-1.90 – 0.26) | 0.20 |
| Chloral Hydrate vs. Slice | 1 | 0.004 | 0.02 (-0.56 – 0.60) | 0.95 |
| Chloral Hydrate vs. Urethane | 1 | 3.129 | -1.67 (-3.00 – -0.34) | 0.08 |
| Isoflurane vs. Slice | 1 | 1.831 | 0.84 (-0.13 – 1.81) | 0.18 |
| Isoflurane vs. Urethane | 1 | 0.609 | -0.85 (-2.96 – 1.26) | 0.44 |
| Slice vs. Urethane | 1 | 3.297 | -1.69 (-2.87 – -0.51) | 0.07 |
| **Post-hoc pairwise comparisons of firing rates based on main effect of genotype/disease model** |  |  |  |  |
| **Comparison** | **DF** | **Chi-Square (Wald Test)** | **Effect Size (95% CI)** | **p-value** |
| WT vs. DBH-Cre | 1 | 1.846 | -0.60 (-1.16 – -0.04) | 0.17 |
| WT vs. TH-Cre | 1 | 2.585 | -0.68 (-1.19 – -0.17) | 0.11 |
| WT vs. VGAT-Cre | 1 | 11.376 | 0.51 (0.12 – 0.90) | 0.0007 |
| WT vs. APP/PS1 | 1 | 0.192 | 0.05 (-0.17 – 0.27) | 0.66 |
| WT vs. DSP-4 | 1 | 5.092 | -0.80 (-1.22 – -0.38) | 0.0249 |
| WT vs. Dyskinetic | 1 | 1.678 | -0.29 (-0.62 – 0.04) | 0.20 |
| WT vs. Parkinsonian | 1 | 0.089 | 0.04 (-0.19 – 0.27) | 0.77 |
| **Post-hoc pairwise comparisons of firing rates based on main effect of age** |  |  |  |  |
| **Comparison** | **DF** | **Chi-Square (Wald Test)** | **Effect Size (95% CI)** | **p-value** |
| Adolescent vs. Adult | 1 | 13.92 | -0.40 (-0.65 – -0.15) | 0.0002 |
| Adolescent vs. Aged | 1 | 16.818 | -0.69 (-1.04 – -0.34) | <0.0001 |
| Adult vs. Aged | 1 | 4.471 | -0.29 (-0.51 – -0.07) | 0.0345 |

**(see excel file for Table S4)**

**Table S4.** Detailed results of the chi-squared statistical test comparing proportion of units in each return map type against the population.

**Movie S1. A rotational view of the manifold.**

A rotational view of the first, second, and third PCs for each ISI histogram (N=1,708 neurons). These are shown over a surface fit to the points (i.e., a manifold).

**Movie S2. A rotational view of points in different areas of the manifold.**

Points from each area are plotted in different colors. The points in each area were randomly selected.

**Movie S3. A rotational view of points in the core of the manifold.**

Points from different areas of the manifold are shown with the PC3 axis truncated to -0.2 to +0.2.

**Movie S4. A rotational view of the points from aged animals in the core of the manifold.**

The ISI distributions of neurons recorded in aged animals are localized to one side of the core.

**Movie S5. A rotational view of the manifold showing points for different genotypes.**

Points for different genotypes are plotted in different colroes (as in Fig. 4J and Fig. 4K). The PC scores for ISI distributions of neurons recorded in VGAT-Cre animals (orange points) are skewed toward one area of the manifold compared to the PC scores for other genotypes.

## Notes

### Summary of Updates

Stylistic shortening of abstract, introduction, and discussion Stylistic merging of multiple figures (and tables) into single figures with many panels (and longer tables) to reduce the number of figures. Reduction of references to match journal limits, in many cases by switching to citation of a review article. Increased discussion on lab-specific effects and the addition of a quantification of across lab variability. The analysis of firing patterns (inter-spike intervals, ISIs) is now done differently and visualized differently, but without any change to the main conclusions (i.e., that firing patterns are diverse and some factors are associated with specific firing patterns). Previously, Figure 6 in our bioRxiv paper showed a 3D scatter plot of the first three Principal Components after doing PCA of the ISI distributions. In that former analysis, we clustered those points (k-means) to find different ISI distributions in the dataset. We then assessed whether different factors (E.g., rats, males, slice recording, etc.) were more predominant in any cluster by comparing the proportion different factors associated with the neurons in each cluster. The problem with this approach is that defining clusters is somewhat arbitrary. We have now embraced this non-clusterable feature of the data and not done any clustering. Instead, we fit a surface (i.e., a manifold) to the 3D scatter plot and show that firing patterns are diverse across different areas of this low-dimensional manifold (Figure 3 and Supplementary Movies 1, 2, and 3). We see some factors are associated with specific areas of the manifold and thus with specific firing patterns (Figure 4 and Supplementary Movies 4, and 5). Thus, rather than carving up the data into artificial clusters and asking what proportion of each cluster is a specific species, or sex, etc., we instead show the low-dimensional spectrum of firing patterns and assess where different factors like species or sex lay along that spectrum.

